# Evolutionary Analysis of Candidate Non-Coding Elements Regulating Neurodevelopmental Genes in Vertebrates

**DOI:** 10.1101/150482

**Authors:** Francisco J. Novo

## Abstract

Many non-coding regulatory elements conserved in vertebrates regulate the expression of genes involved in development and play an important role in the evolution of morphology through the rewiring of developmental gene networks. Available biological datasets allow the identification of non-coding regulatory elements with high confidence; furthermore, chromatin conformation data can be used to confirm enhancer-promoter interactions in specific tissue types and developmental stages. We have devised an analysis pipeline that integrates datasets about gene expression, enhancer activity, chromatin accessibility, epigenetic marks, and Hi-C contact frequencies in various brain tissues and developmental stages, leading to the identification of eight non-coding elements that might regulate the expression of three genes with important roles in brain development in vertebrates. We have then performed comparative sequence and microsynteny analyses in order to reconstruct the evolutionary history of the regulatory landscape around these genes; we observe a general pattern of ancient regulatory elements conserved across most vertebrate lineages, together with younger elements that appear to be mammal and primate innovations. This preprint has been reviewed and recommended by Peer Community In Evolutionary Biology (http://dx.doi.org/10.24072/pci.evolbiol.100035)

## INTRODUCTION

Large-scale genomic projects (Andersson et al. 2014; Kundaje et al. 2015) have identified thousands of regulatory regions in the human genome that behave as enhancers or silencers of gene expression and are defined by specific epigenetic modifications, chromatin accessibility and occupancy by transcription factors. Many of these regulatory elements are highly conserved in the genomes of various animal groups and regulate the expression of genes involved in developmental pathways. Several studies have shown that conserved non-coding elements (CNEs) underwent an expansion in early vertebrate evolution, particularly during the transition from Agnatha to Gnathostomata (McEwen et al. 2009).

Evolutionary changes in CNEs mirror macroevolutionary trends in morphology and anatomy, as shown in a study by Lowe et al. in which they identified several million conserved non-exonic enhancer elements and determined when such elements became regulatory over the last 600 million years (My) and which genes they regulate (Lowe et al. 2011). Notably, they found that the most ancient set of CNEs (those that became active between 600 and 300 My ago) regulate the activity of transcription factor and developmental genes. This confirmed previous suggestions that extensive rewiring of developmental networks took place at the time of diversification of animal body plans (Vavouri and Lehner 2009; Pauls et al. 2015). In fact, the same basic set of regulatory genes controls development in different animal groups; therefore, different animal morphologies can be explained by the re-deployment of the same genes in various alternative regulatory circuits, leading to gene-regulatory networks of different topologies (Erwin and Davidson 2009). In this regard, evolution of non-coding regulatory DNA is a major contributor to the evolution of form (Carroll 2005).

The role of CNEs in morphological evolution is also relevant in the context of the evolution of the brain, particularly the neocortex (Rakic 2009; Bae et al. 2015). In a landmark study, Pennacchio et al. (Pennacchio et al. 2006) characterized the *in vivo* enhancer activity of a large group of non-coding elements that are conserved in human-pufferfish or in human-mouse-rat; the majority of these elements directed gene expression in various regions of the developing nervous system. Wenger et al. (Wenger et al. 2013) used p300 ChIP-seq to identify several thousand enhancers active at embryonic day E14.5 in mouse dorsal cerebral wall. Reilly et al. (Reilly et al. 2015) have recently identified thousands of regulatory elements involved in the formation of the neocortex during the first weeks of embryonic development in the human lineage but not in macaque or mouse. A remarkable effort by Boyd et al. (Boyd et al. 2015) showed that insertion in the mouse genome of a human CNE regulating the expression of a neurodevelopmental gene led to a 12% increase in brain size in mice, whereas the chimpanzee homologous element did not.

Current understanding of the structure and function of regulatory elements within the 3D organization of chromatin shows that promoters frequently make contact with distal elements located tens or hundreds of kb away (McLean et al. 2010; Wenger et al. 2013; Shlyueva et al. 2014; Smemo et al. 2014) if they are in the same topologically associating domain (TAD) (Dekker and Mirny 2016; Dixon et al. 2016). In this work we have used existing datasets providing genomic features typical of regulatory elements (chromatin accessibility, epigenetic marks, binding of transcription factors) and merged them with previously defined catalogues of brain-specific enhancer elements in mouse and human (Pennacchio et al. 2006; Wenger et al. 2013; Andersson et al. 2014). Incorporating into our analysis pipeline information about chromatin contacts in human developing brain (Won et al. 2016) allowed us to make high-confidence predictions about enhancer-promoter interaction and identify eight putative regulatory elements for three important neurodevelopmental genes. Comparative analyses of these elements in vertebrate genomes enabled us to reconstruct the evolutionary history of the regulatory landscape around these genes.

## RESULTS

### 1. Identification and characterization of candidate regulatory elements

We initially performed various intersections of four primary datasets (2,019 predefined brain-specific FANTOM5 enhancers; 6,340 mouse telencephalon p300 peaks; 1,339 Vista enhancers; 36,857 ancient CNEEs) in order to find genomic regions with high probability of behaving as deeply conserved brain-specific developmental regulatory elements. For the most promising candidates passing these filters, we manually inspected TF binding data and epigenomic marks to support their function as active enhancers in brain cells/tissues.

For every candidate regulatory element studied, the next crucial step was to predict which of the genes in the vicinity was more likely to be regulated by the element. To this end, we first obtained a comprehensive view of all protein-coding and non-coding genes annotated in the region generated by the GENCODE project and mapped to GRCh37/hg19 (UCSC Genome Browser GENCODE Genes track, comprehensive set version 24lift37). We looked at gene expression in GTEx and Brainspan, and then selected those genes with preferential expression in brain. Next, we searched GTEx for eQTLs near candidate regulatory elements in order to predict which genes might be regulated by variants in the same region. We also searched for SNPs in the vicinity that had been previously associated with brain-related phenotypes in GWAS studies. Finally, and most importantly, we investigated the frequency of physical contacts between the candidate regulatory element and the promoters of the genes in the region, using Hi-C data from human fetal brain and comparing it to Hi-C data from other cells/tissues.

The result of this analysis pipeline was a list of candidate Brain-specific Regulatory Elements (BREs) in humans, together with the gene(s) predicted to be regulated by each element. We focused our analysis on BREs regulating protein coding genes *TBR1*, *EMX2* and *LMO4* because they code for transcription factors known to be involved in brain development. Since enhancer-gene associations can be established with high confidence thanks to Hi-C evidence, we will present the results organized around these three genes. Table 1 shows a summary of the regulatory elements identified in this study.

**Table 1.** Brain-specific regulatory elements (BREs) identified and analysed in this work. For every BRE, it shows GRCh37/hg19 coordinates, and the HGNC name and the coordinates of the genes regulated by them (coordinates of the longest genomic-spanning variant in GENCODE v24lift37, comprehensive set).

| Regulatory element ID | BRE coordinates (GRCh37/hg19) | Regulated gene | Gene coordinates (GRCh37/hg19) |
| --- | --- | --- | --- |
| BRE1 | chr2:162,095,064-162,095,312 | *TBR1* | chr2:162,272,605-162,282,381 |
| BRE2 | chr10:119,293,007-119,295,403 | *EMX2* | chr10:119,301,955-119,309,056 |
| BRE3 | chr10:119,310,654-119,312,169 | *EMX2* |  |
| BRE4 | chr10:119,432,340-119,433,475 | *EMX2* |  |
| BRE5 | chr10:119,786,696-119,790,021 | *EMX2* |  |
| BRE6 | chr1:87,600,259-87,603,242 | *LMO4* | chr1:87,794,151-87,814,606 |
| BRE7 | chr1:87,821,614-87,822,820 | *LMO4* |  |
| BRE8 | chr1:88,758,718-88,759,914 | *LMO4* |  |

### 2. *TBR1* (ENSG00000136535)

The first regulatory element identified (BRE1) is a 248 nt fragment located 2.4 kb from the 3’-end of the longest *TANK* (Homo sapiens TRAF family member-associated NFKB activator) variant (Figure 1). It is embedded in a region with Roadmap epigenetic marks typical of active Transcription Start Site (TSS, red) and binding of several TFs including RNApol2. However, there is no promoter overlapping this region or anywhere nearby: the closest promoters belong to ncRNA *AC009299.3 (LINC01806)* located 6 kb downstream and to ncRNA *AC009299.3* located 7.6 kb downstream. None of these two genes is expressed in brain: according to GTEx, *AC009299.3* is expressed almost exclusively in thyroid, and *AC009299.2* only shows expression in testis.

**Figure 1.**
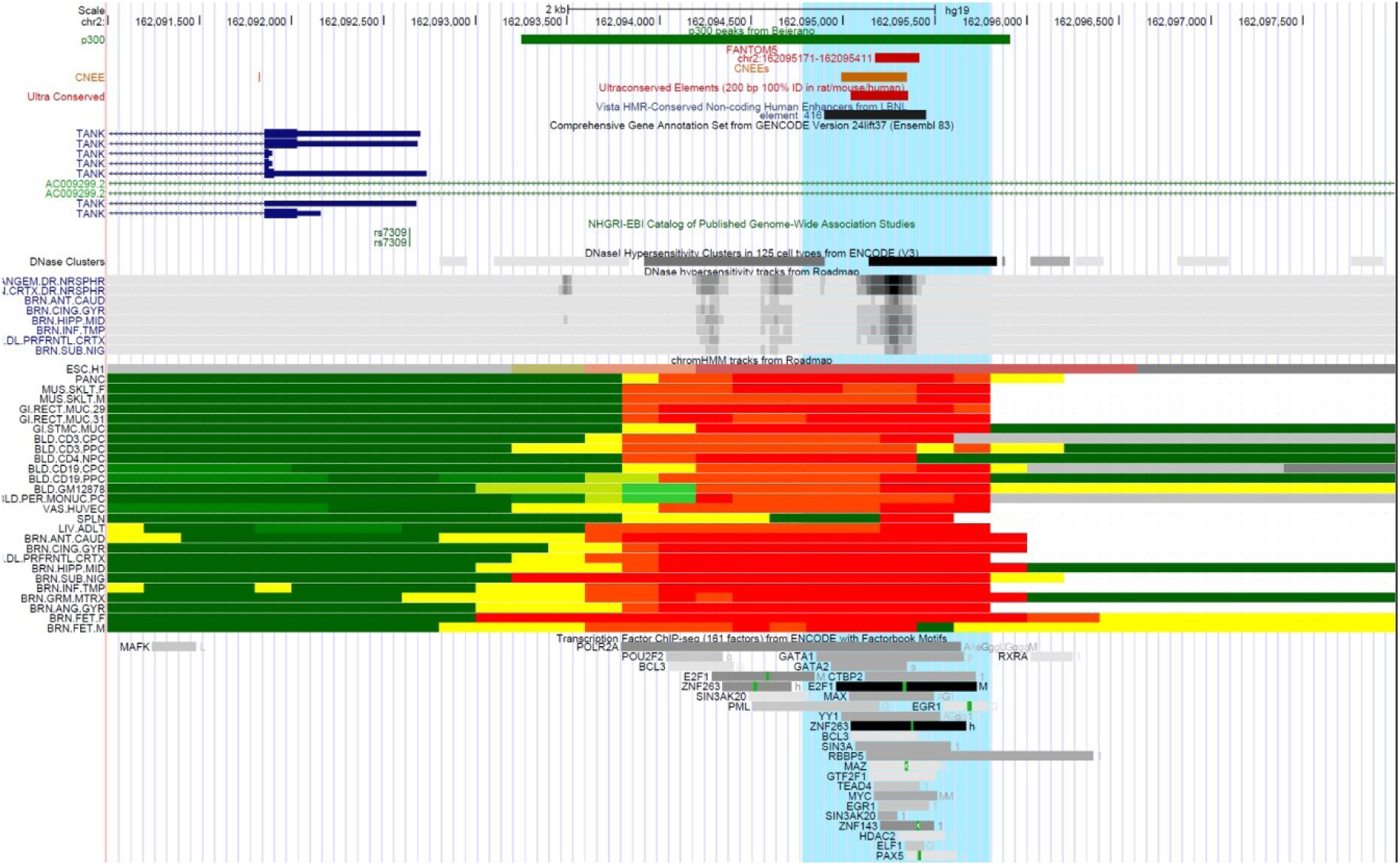
Genomic landscape of BRE1 (located within the area highlighted in light blue). UCSC tracks (from top to bottom): p300 peaks in mouse telencephalon (green), FANTOM5 predefined brain-specific enhancers (red), conserved non-exonic enhancers (brown), ultraconserved elements (red), Vista enhancers (black), GWAS SNPs (green), Comprehensive GENCODE annotations, GWAS SNPs (green), DHS from ENCODE, DHS from Roadmap, ChromHMM predictions from Roadmap, and Transcription Factor ChIP-seq from ENCODE. These tracks can be viewed at this UCSC Genome browser session: http://genome.ucsc.edu/cgi-bin/hgTracks?hgS_doOtherUser=submit&hgS_otherUserName=fnovo&hgS_otherUserSessionName=BRE1

The epigenetic marks flanking the element are typical of active enhancer (yellow in chromHMM tracks). These marks are present both in brain and non-brain tissues from Roadmap (see methods for tissues included), suggesting that this element can regulate target genes in different tissues. Notably, BRE1 overlaps one p300 peak, one FANTOM5 predefined brain-specific enhancer, and one Vista element (element 416) that is very active in the forebrain of E11.5 mouse embryos. All these lines of evidence support a clear role for BRE1 as a regulatory element active in developing brain.

In order to predict which of the neighbouring genes are regulated by this element in brain tissues, we looked at virtual 4C profiles derived from Hi-C studies. Although BRE1 is 2.4 kb from the 3’-end of *TANK*, we found no contacts between this element and the promoter of *TANK* in fetal brain samples from the cortical plate and germinal zone (Figure 2). In fact, the two main peaks of contacts with BRE1 in these tissues encompass the promoters of ncRNAs *AC009299.3* and *AC009299.2* (proximal peak, Figure 2) and the promoter of *TBR1* (177 kb downstream, distal peak to the right in Figure 2). In GM12878 and IMR90 cell lines (which are not of neural origin) these contacts are absent (not shown). These findings suggest that BRE1 might function as a regulatory element responsible for the expression of ncRNAs *AC009299.3* and *AC009299.2* in non-brain tissues and of *TBR1* in developing brain. Of note, *TBR1* is clearly involved in neurodevelopmental processes: it shows brain-specific expression in cortex according to GTEx and in neocortex during early brain development according to Brainspan. It has been previously shown that it is expressed in the pallium during brain development in vertebrates from zebrafish to mouse (Rakic 2009; Wenger et al. 2013). To gain further insights into *TBR1* regulation in brain, we explored which regions make contact with its promoter using Hi-C data and confirmed our initial findings: although no distal regions contact this promoter in GM12878 and IMR90 cells, contacts with BRE1 are high in fetal brain tissues (Supplementary Figure S1).

**Figure 2.**
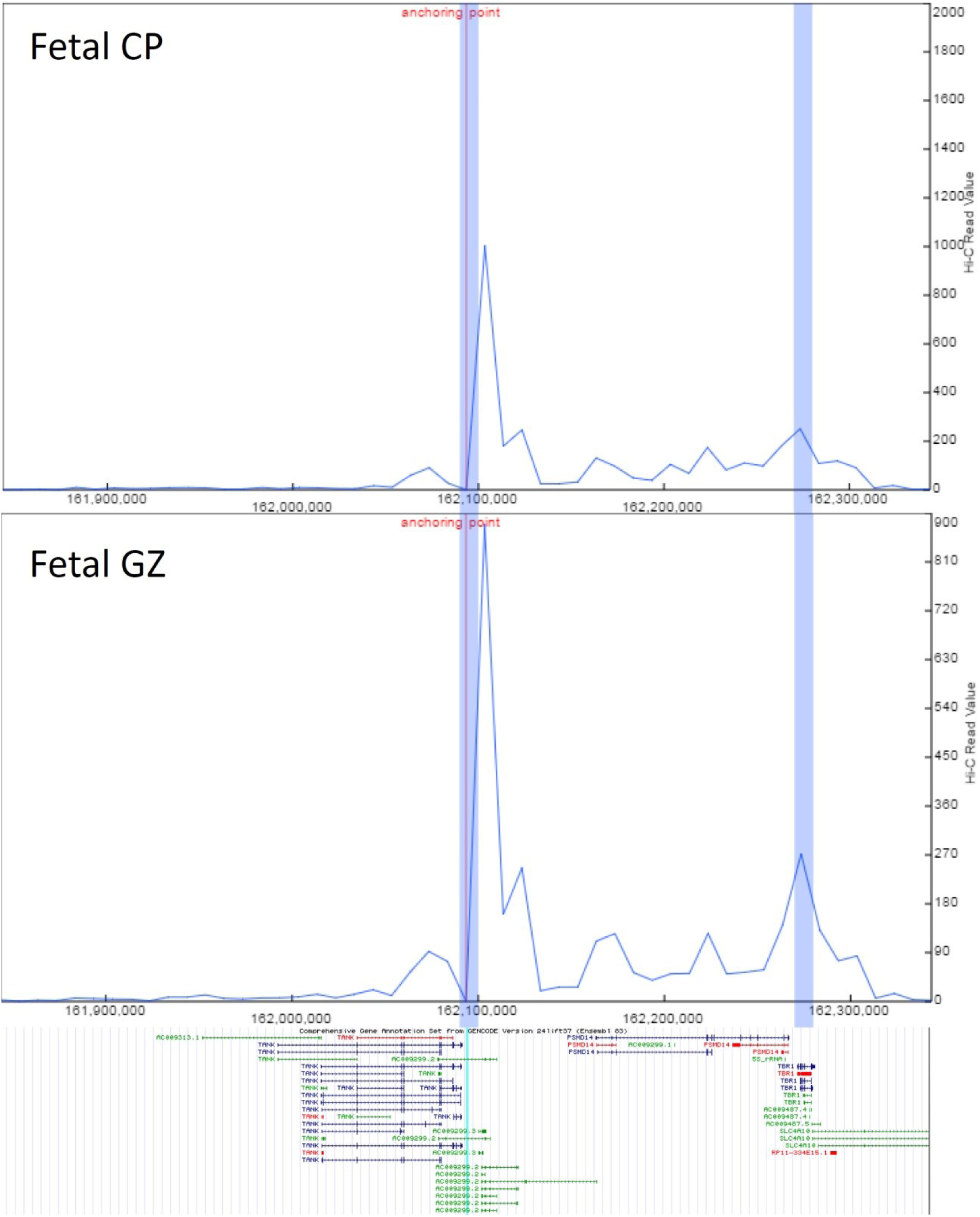
Virtual 4C plots using Hi-C data from fetal brain cortical plate (Fetal CP) and germinal zone (Fetal GZ) from (Won et al. 2016). In each plot, the anchoring point (red line) marks the position of BRE1 (thin light blue line in the bottom panel). There is a peak of contacts between BRE1 and the promoters of ncRNAs *AC009299.3* and *AC009299.2* (proximal peak highlighted in blue in the middle) and the promoter of *TBR1* (distal peak highlighted in blue to the right). The bottom panel shows the GENCODE gene annotation comprehensive set, version 24lift37. No contacts are evident between BRE1 and the promoter of *TANK* (to the left). Contacts (Y-axis) are shown as RPKMs.

We also searched GTEx for overlapping eQTLs that could shed some information on potential genes regulated by this element. Although some eQTLs affect the expression of *TANK* and AC009299.2 in other tissues, no eQTLs are annotated in brain samples. We also note the presence of a GWAS SNP nearby (rs7309, right at the 3’-end of *TANK*) associated with educational attainment (Okbay et al. 2016). This SNP is in a linkage disequilibrium block which ends abruptly at rs11678980, close to the start of AC009299.3.

As shown in Figure 1, BRE1 overlaps ultraconserved element uce-6261 located at chr2:162297586-162297897 in assembly hg16 (uc.88 in (Bejerano 2004)) which corresponds to chr2:162095042-162095353 in GRCh37/hg19. There is also an overlapping ancient CNEE, suggesting that this is a region of deep conservation. This led us to perform blast searches on representative genomes from Porifera and Ctenophora to primates. COGE blastn wtih the human sequence as query detected significant hits (E-value < 1e-06) in elephant shark, spotted gar, zebrafish, fugu, coelacanth, mouse, and human; no hits were detected in lamprey or earlier species. This suggests that BRE1 is a vertebrate innovation appearing in Gnathostomes. The hit in elephant shark covers the first 120 nt of the query and it is expanded in Osteichthyes, where it covers almost the entire query. The hit is slightly shorter in xenopus, but in chicken and mammals it covers again the whole query sequence (Supplementary Figure S2).

We then studied the microsynteny of the region surrounding the BRE1 element (Figure 3). We observed that coelacanth, spotted gar and elephant shark have orthologs for *TANK*, *PSDM14* and *TBR1* in the same order and orientation than mammals, with another gene *RBMS1* upstream from *TANK* in the opposite orientation (the probability of finding this arrangement of 4 random genes is 1/384 or 0.003). Lamprey also has orthologs for *RBMS1* (PMZ_0005867-RA located at GL482213:117724-268242 in WUGSC 7.0/petMar2 genome assembly), *PSMD14* (PMZ_0005863-RA at GL482213:59613-84084) and *TBR1* (PMZ_0005862-RA at GL482213:17651-47396), but not for *TANK*. However, the *RBMS1* lamprey ortholog (PMZ_0005867-RA) is transcribed in the same orientation as the other genes in the region, contrary to the situation in all the other species. This suggests that an inversion took place in the transition from lamprey to elephant shark, which could have brought the sequence of BRE1 close to *TBR1*. In fact, there is another lamprey gene annotated between the orthologs for *RBMS1* and *PSMD14* (PMZ_0005864-RA at GL482213:91955-99066); blastp shows that it is the ortholog of mammalian *AZI2*. Interestingly, *TANK* probably originated by a duplication of the *AZI2* gene according to GenTree.

**Figure 3.**
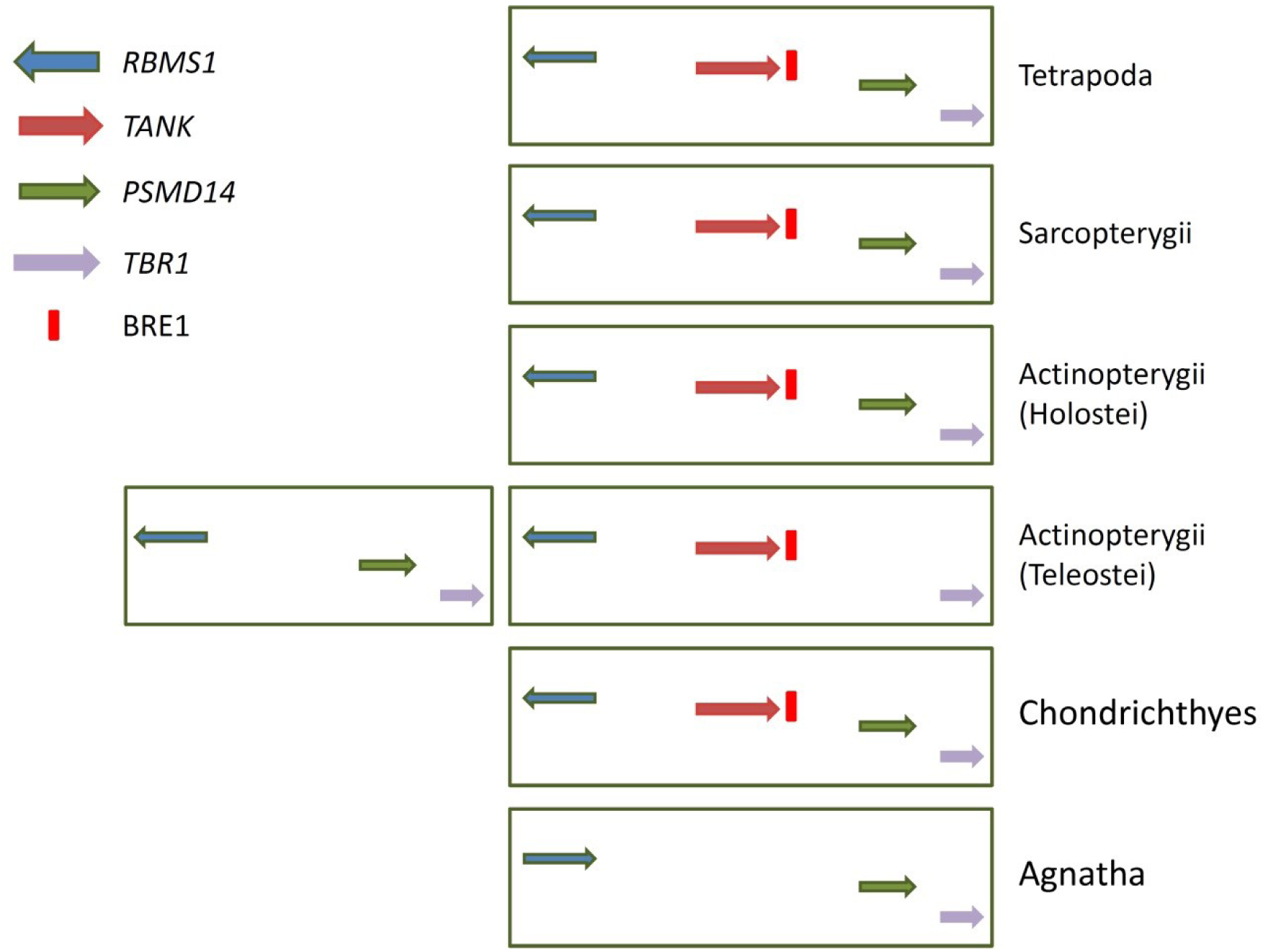
Schematic representation of the evolution of the BRE1 element and microsynteny of the region. Coloured arrows represent the orthologs of the four genes in the region; the red segment represents the BRE1 element. Sarcopterygians are represented by the coelacanth, Holostei by spotted gar, Teleostei by zebrafish, fugu, tetraodon and stickleback, Chondrichthyes by elephant shark, and Agnatha by lamprey. The additional left rectangle in Teleostei indicates the presence of a duplication in a different chromosome.

We looked at the presence of repeats that could explain the origin of this region by transposition-mediated transduction, but there is no obvious repeat to account for such an event: the BRE1 element is flanked by a SINE2-1_CM and a REP1_CM repeat in elephant shark, but no traces of the region (including these repeats) are detected by BLAT in the genomes of the lamprey or branchiostoma. Taken together, these data suggest that an inversion of the *RBMS1* ortholog during the transition from lamprey to gnathostomes, concurrent with the duplication of the *AZI2* lamprey ortholog, is the most likely cause leading to the appearance of *TANK* in gnathostomes; the BRE1 element would have been generated during the same process.

Some members of infraclass Teleostei (zebrafish, fugu, tetraodon and stickleback) have a *TANK* ortholog but no *PSMD14* ortholog in the region. The absence of the latter leaves a gap between the BRE1 element and *TBR1*. However, in these species there is a *PSMD14* ortholog in a different chromosome, very close to another *TBR1* ortholog (*Tbr1a*) but without any trace of the BRE1 element sequence. As these changes are not seen in the genome of spotted gar (*Lepisosteus oculatus*, infraclass Holostei), this is probably the result of the whole-genome duplication event that took place in Teleostei (Inoue et al. 2015). After duplication, the new copies of these genes underwent different evolutionary pathways: the ortholog for *PSMD14* was lost in the copy including *TANK* and the BRE1 element, whereas the other copy lost the *TANK* ortholog and the BRE1 element. These configurations are summarized in Figure 3.

In order to further clarify the origin and potential functional significance of the BRE1 element we searched for its presence in transcribed sequences, using blastn on EST and RefSeq RNA databases. We found a significant match with the first 250 nt of JW604660, a mRNA sequence (813 total length) from *Fundulus grandis* (order Cyprinodontiformes); with nucleotides 215 to 450 of mRNA FE203853.1 from the brain of *Dissostichus mawsoni* (antarctic toothfish); with part of the 3’-UTR of *TANK* mRNAs of *Stegastes partitus* and *Notothenia coriiceps* (both superorder Acanthopterygii); with the final exon of a ncRNA variant of *TANK* in *Esox Lucius* (superorder Proacanthopterygii); and with part of a mRNA from donkey coding for cell wall protein gp1-like *LOC106843135*. These findings are presented schematically in Supplementary Figure S3. In summary, it appears the in Acanthopterygii (particularly in order Perciformes) the enhancer was either part of the 3’-UTR of a long *TANK* mRNA or part of other brain-expressed mRNAs. In order Cypriniformes (as seen in zebrafish) it became separated from *TANK*. In donkey it became engulfed by another gene, apparently unique to this species. No traces of expression of the BRE1 element were found in any other species.

## 3. *EMX2* (ENSG00000170370)

One of the regions identified in our initial screening, showing very good epigenetic marks in chromHMM Roadmap tracks, is located 1.6 kb from the 3’-end of *EMX2* (empty spiracles homeobox 2). This gene is expressed in brain although it is not brain-specific according to GTEx and Human Protein Atlas data (ovary and uterus being the organs with highest expression). However, it is expressed in early embryonic hippocampus (archipallium) according to Brainspan, it is a negative regulator of *SOX2* in neural stem cells in telencephalon, and it may function in combination with *OTX1/2* to specify cell fates and to generate the boundary between the roof and archipallium in the developing brain (Cecchi et al. 2000; Rakic 2009; Mariani et al. 2012).

The involvement of *EMX2* in brain development led us to search for additional regions that might regulate its expression. First, we found some GTEx eQTLs for *EMX2* located between 60-100 kb upstream of its TSS. In addition, we analysed Hi-C data from fetal brain in a 1 Mb area surrounding the gene, in order to identify regions that contact its promoter. As shown in Figure 4, several regions display significant contacts by virtual 4C analysis. Further inspection of Roadmap epigenetic marks in those regions led us to select four potential regulatory elements of *EMX2*, including the one initially found: BRE2, BRE3, BRE4 and BRE5.

**Figure 4.**
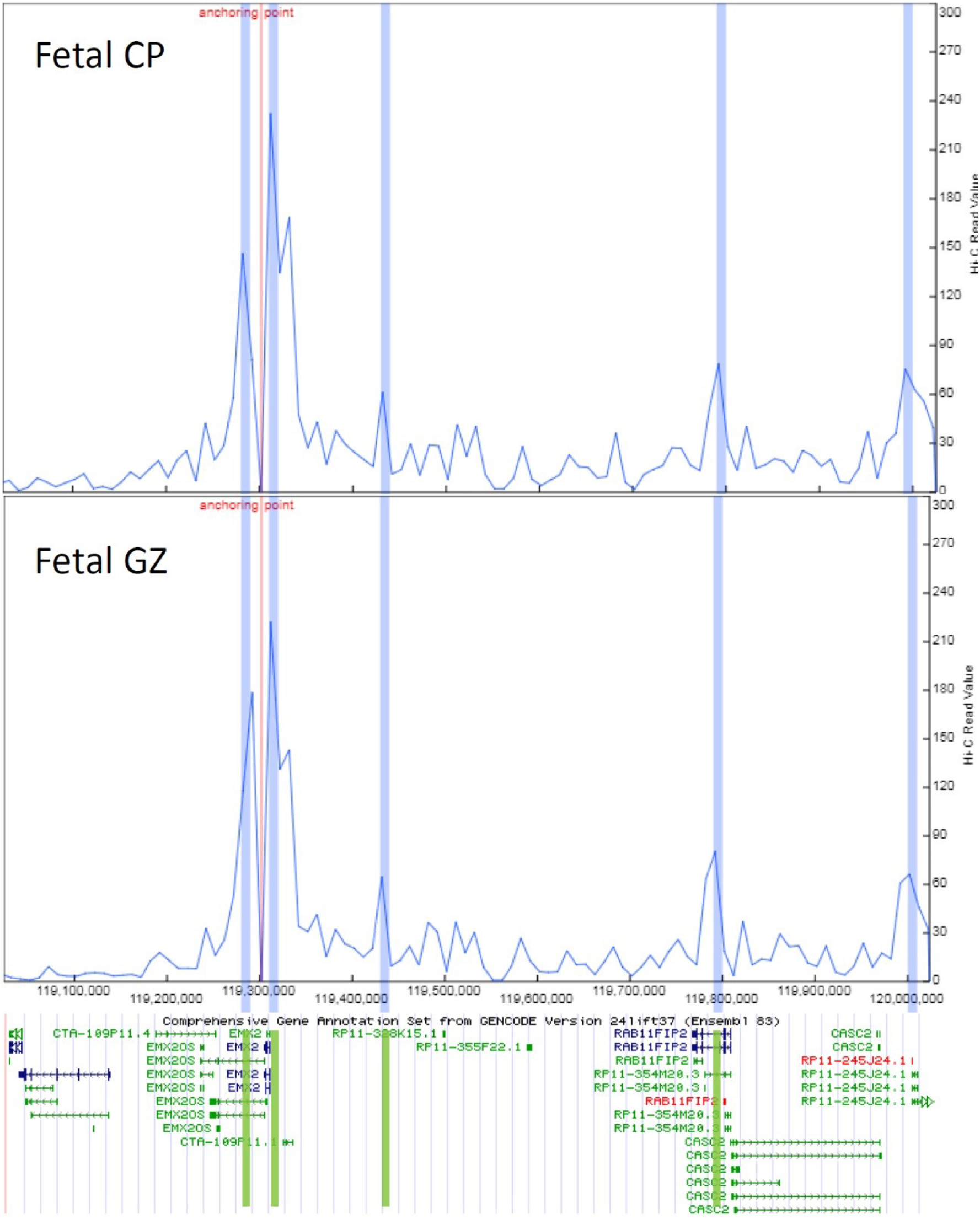
Virtual 4C plots using Hi-C data from fetal brain cortical plate (Fetal CP) and germinal zone (Fetal GZ) from (Won et al. 2016). In each plot, the anchoring point (red line) marks the position of the *EMX2* promoter. The bottom panel shows the GENCODE gene annotation comprehensive set, version 24lift37. Contacts (Y-axis) are shown as RPKMs. Five peaks of contacts are clearly visible (regions highlighted in blue); the first four (from left to right) correspond to BRE2, BRE3, BRE4, and BRE5, respectively (highlighted in green in the bottom panel).

### 3.1. BRE2

(chr10:119,293,007-119,295,403) is a regulatory region located 6.6 kb upstream the TSS of *EMX2*, so it is not part of its core promoter. The element does not overlap any Vista enhancer, FANTOM5 predefined brain-specific enhancer or p300 peak from mouse telencephalon. Roadmap epigenetic marks classify it as active enhancer only in brain tissues, liver, spleen, pancreas and gastrointestinal mucosa, but not in blood cells. Some TFs bind to this region in several tissues according to ENCODE data. Conservation analysis with COGE blast found no traces of this sequence in lamprey, shark, osteichthyans,coelacanth, xenopus or platypus; there is a short hit in opossum, with an additional hit in mouse. The entire first half of the region is present in small-eared galago (*Otolemur garnettii*); in marmoset (*Callithrix jacchus*) and apes the whole BRE2 element is perfectly conserved (Supplementary Figure S4).

The 4.3 kb genomic region containing BRE2 is flanked by two MIR SINE repeats in the human genome (MIRc on one side and MIRb on the other). The same happens in mouse, where the syntenic 4 kb region is flanked by MIR repeats. This raises the possibility that this regulatory region was added to mammalian genomes by a retrotransposon-mediated transduction event.

In order to find any potential expressed sequences containing this region, we performed a blastn search on RefSeq RNA in Vertebrata (taxid:7742), finding two significant hits: the first (E-value=0.0) shows the whole query in the final half of XR_002082680.1 from *Panthera pardus*; the second hit (E-value=1e-130) shows the first half of the query covering almost the entire length (except for the first 213 nt) of XR_782286.2 from the mole rat *Fukomys damarensis*. Both records are unannotated ncRNAs of the type LOCxxxxxxxxx, so the significance of this finding is uncertain.

### 3.2. BRE3

(chr10:119,310,654-119,312,169) is a regulatory region located 1.6 kb from the 3’-end of *EMX2*. It overlaps one Vista enhancer very active in midbrain (element 935), and one FANTOM5 predefined brain-specific enhancer. The region corresponding to the Vista enhancer also overlaps one ancient NCEE. It shows Roadmap epigenetic marks of active enhancer in several brain tissues, liver, spleen, pancreas and skeletal muscle, as well as binding of several TFs according to ENCODE.

Conservation analysis with COGE blast shows a 300 nt hit (corresponding to the region of the Vista enhancer) in elephant shark, coelacanth, xenopus, chicken and opossum; nothing was detected in lamprey or earlier genomes. An additional hit appears in mouse which is extended in galago and mouse lemur; in the marmoset and later primates there is a single hit covering the entire query with almost perfect similarity (Supplementary Figure S5).

There is a MER91A DNA-transposon 3.7 kb away from one end of this element in the human genome, but the first transposon on the other side is 12 kb away. In the oldest primate analysed (*Otolemur garnettii*) there is a similar situation, with a MER91A repeat on one side of the syntenic region. However, no recognizable transposons are found in the vicinity of the syntenic region in mouse, so it is unclear whether this regulatory element was added to primate genomes via a transposon-mediated mechanism.

In order to find any potential expressed sequences containing this region, we did a blastn search on RefSeq RNA in all species but we only found one hit: the first 450 nt of the query are part of the central region of one predicted *EMX2* mRNA from *Astyanax mexicanus* (XM_007238578.2), not well characterized.

### 3.3. BRE4

(chr10:119,432,340-119,433,475) is a 1.1 kb element overlapping a clear peak of Hi-C contacts with the promoter of *EMX2*, located 125 kb away. It does not overlap any Vista enhancer, FANTOM5 predefined brain-specific enhancer or p300 peak, but it has strong active enhancer marks exclusively in fetal brain according to Roadmap data. ENCODE ChIP-seq data shows binding of EBF1 in the only cell line in which it was assayed.

COGE blast with an internal fragment of 847 nt (chr10:119,432,372-119,433,218) only found hits in primates (covering most of the query) from *Otolemur garnettii* to *Pan troglodytes*. Additionally, a short internal fragment of 77 nt was found in mouse, in a syntenic region 120 kb from the 3’-end of *Emx2*, where it is flanked by a MIRb SINE element only on one side. In the human genome this element is flanked by MIR3 and MIRb SINE elements, and in the oldest primate analysed *(Otolemur garnettii)* it is also flanked by MIR SINE elements on either side (MIRb and MIRc respectively), so it might have been the result of retrotransposon-mediated transduction. This regulatory element is not expressed, as blast searches on RefSeq RNA (in all RefSeq) were negative.

### 3.4. BRE5

(chr10:119787196-119788657) is a 1.4 kb regulatory region with strong epigenetic marks of active enhancer almost exclusively in brain tissues according to Roadmap data, and with binding of several TFs in ENCODE cell lines. It does not overlap any Vista enhancer, p300 peak or FANTOM5 predefined enhancer, but Hi-C data in fetal brain shows a strong peak of contacts with the promoter of *EMX2* located 485 kb away. The element lies inside an intron of *RAB11FIP2* (a gene also expressed in brain, although not brain-specific), but it does not seem to contact the promoter of this gene at least in fetal brain Hi-C data.

COGE blast revealed that this region was added in primates, because there are no traces in genomes older than the galago *Otolemur garnettii*, where it is almost completely conserved. It is flanked by a Tigger14a DNA transposon and a MIR SINE on one side, and by an L2c LINE repeat on the other. The syntenic region in *Otolemur* is also flanked by the Tigger14a DNA transposon and a MIRb SINE on one side, so it is likely that it was incorporated into primate genomes by a transposon-mediated event. Searching for expressed sequences in RefSeq RNA (all species) was negative.

Microsynteny analysis of this region of chromosome 10 revealed the presence of *EMX2* flanked by gene *RAB11FIP2* on one side (in the opposite orientation) and genes *PDZD8* and *SLC18A2* on the other side of *EMX2* (Figure 5). This configuration is preserved in the genome of the elephant shark, the coelacanth, marsupials and eutherian mammals. As shown in Figure 5, Teleostei (zebrafish and fugu), xenopus and platypus seem to have lost some of the genes in the region. Although *EMX2* and its closest regulatory element (BRE3) are present from the elephant shark to primates, BRE2 appears in marsupials, BRE4 appears in eutherian mammals and BRE5 is exclusive to primates.

**Figure 5.**
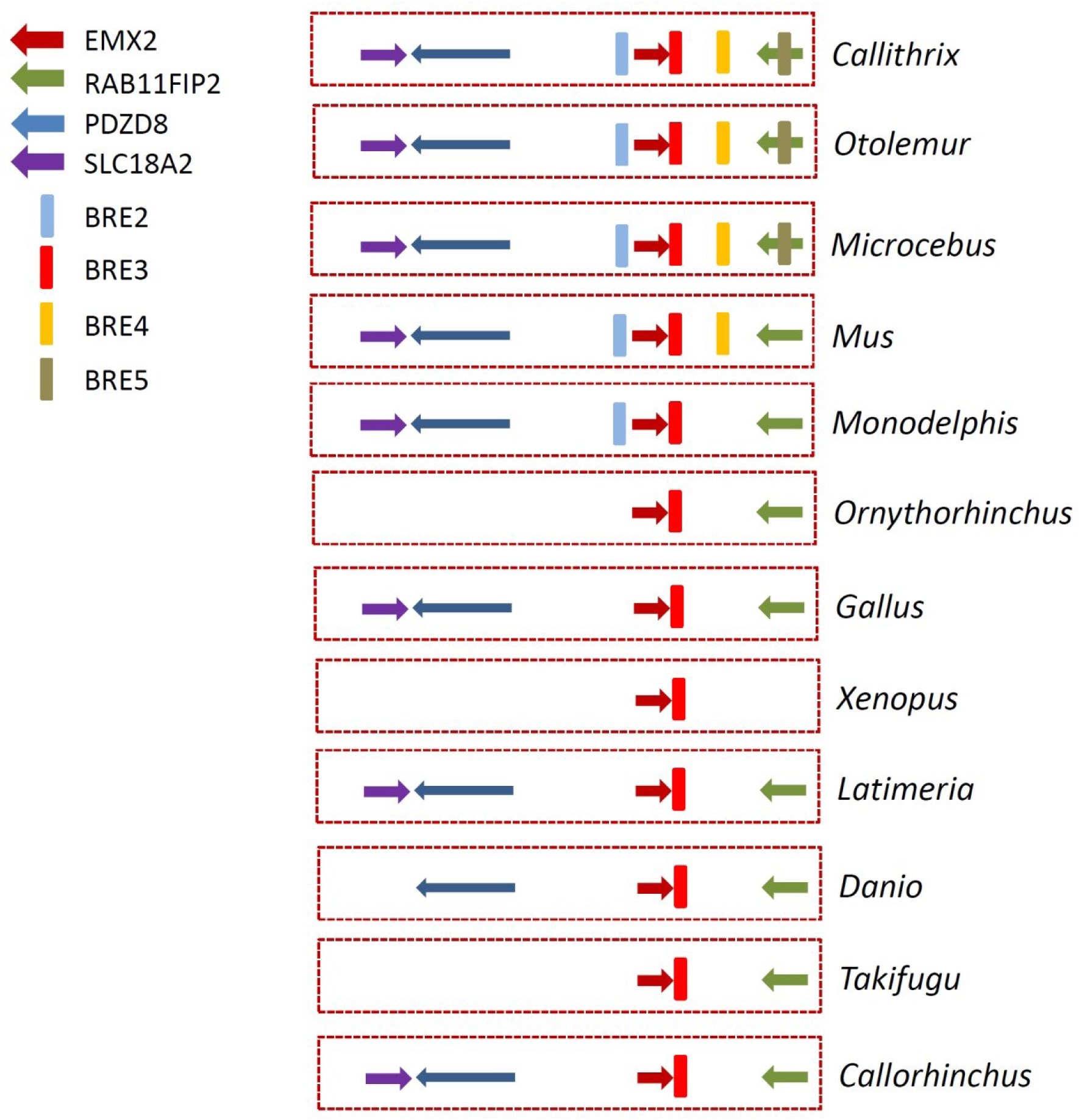
Schematic representation of the evolution of regulatory elements of *EMX2* and microsynteny of the region. Coloured arrows represent the orthologs of the four genes in the region, coloured segments represent the BREs.

## 4. *LMO4* (ENSG00000143013)

Our initial screening also highlighted a good candidate enhancer element located 7.1 kb from the 3’-end of the longest mRNA variant of protein coding gene *LMO4* (LIM domain only 4; the protein encoded by this gene contains two LIM domains). This gene probably originated by duplication from *LMO1* in Eutelostomi, according to GenTree. It shows good brain-specific expression in GTEx and also high expression in fetal brain by BrainSpan. Analysis of *Lmo4* expression during mouse brain development shows increased regionalization in mid-and late-embryonic stages, suggesting a role for this gene in the specification of certain brain regions (Hermanson et al. 1999; Bulchand et al. 2003).

GTEx does not provide information on eQTLs regulating this gene. Several GWAS SNPs in the region are related to neurological phenoptypes: SNPs rs72953990 (associated to human gait problems) and rs35214987 (associated with cognition) are located about 300 kb from *LMO4*; SNPs rs74823926 (associated with cannabis dependence) and rs2179965 (associated with cognitive test measures) are 1 Mb away from *LMO4*. Analysis of Hi-C data from fetal brain, compared to GM12878 and IMR90 cells, identified several regions with increased number of contacts in fetal cortical plate and germinal zone (Supplementary Figure S6). These lines of evidence, together with inspection of Roadmap epigenetic marks, led us to select three potential regulatory regions of *LMO4* (BRE6, BRE7 and BRE8) for further analysis.

### 4.1. BRE6

(chr1:87,600,259-87,603,242) is a 3 kb element located 190 kb upstream of *EMX2* and corresponds to a peak of Hi-C contacts that overlaps *LINC01140*. The proximal promoter of this lncRNA shows marks of active TSS in all tissues, but its expression is low in brain. The BRE6 regulatory element is further downstream towards the 3’-end of *LINC01140* and shows marks of active enhancer only in brain tissues according to Roadmap.

COGE blast found significant hits in coelacanth and xenopus, corresponding to nucleotides 1,900-2,100 of the query. This region of similarity is progressively extended in chicken and mammals, so that in *Otolemur* it encompasses almost the entire BRE6 element. We looked at the microsynteny of the region, where *HS2ST1* is close to *LMO4* (both genes in the same orientation) in all vertebrates. BRE6 is 25-30 kb away from the 3’-end of *HS2ST1* in primates and at a similar distance in mouse, but much further away in opossum. In platypus, the element is inside the *HS2ST1* ortholog, but in chicken, xenopus and coelacanth the distance is similar to that in mammals (Figure 6). These searches also found parts of the future ncRNA *LINC01140* overlapping this enhancer region, which (in addition to behaving as regulatory element of *LMO4)* apparently became part of the final long exon of the short variants of human *LINC01140*. Searches for expressed sequences (blast on RefSeq RNA, all species) only found hits in primates, corresponding to the part of human *LINC01140* that overlaps this region.

**Figure 6.**
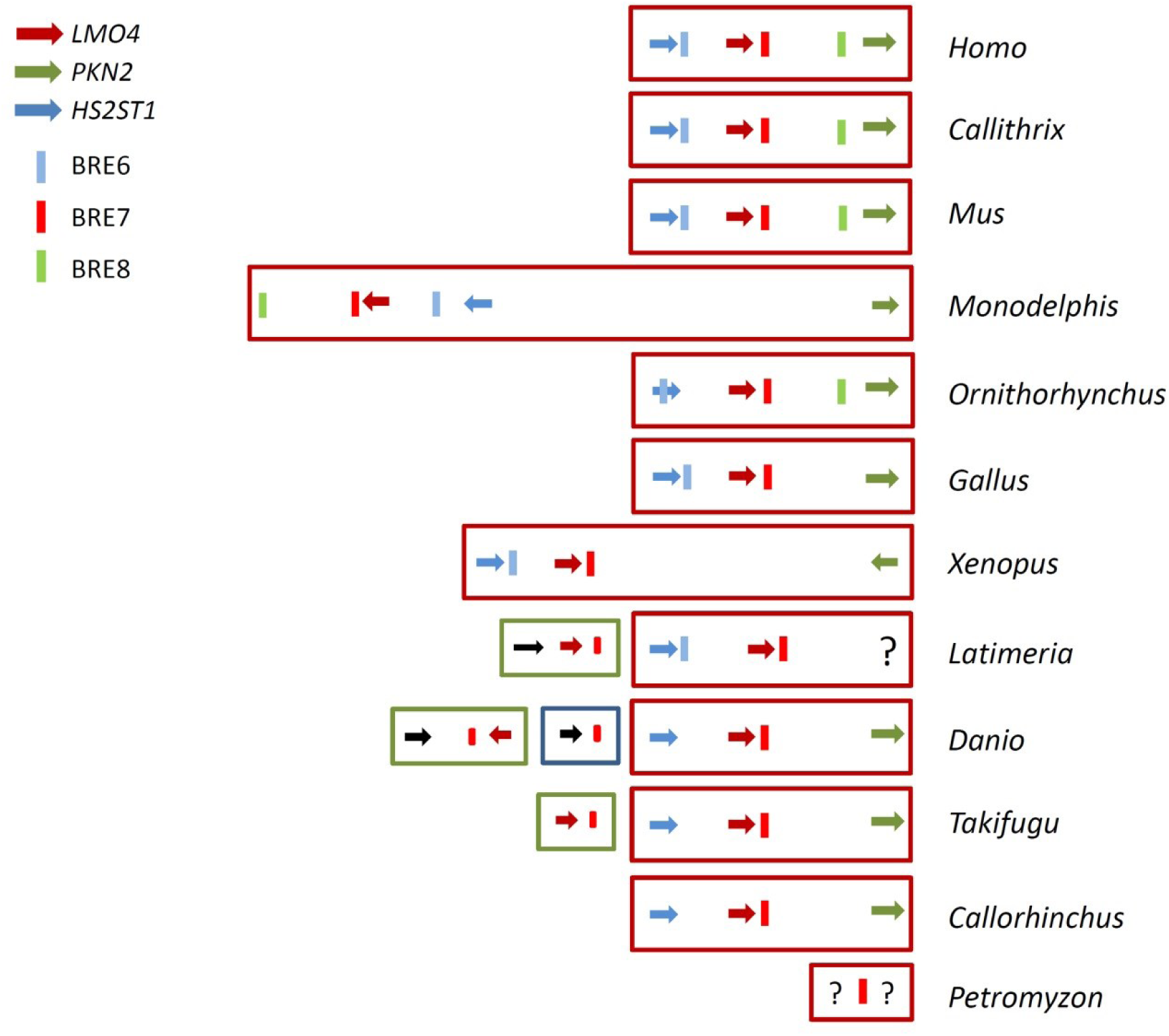
Schematic representation of the evolution of regulatory elements of *LMO4* and microsynteny of the region. Coloured arrows represent the orthologs of the three genes in the region, coloured segments represent the BREs.

The element is next to a MIRb SINE repeat on one side in human, chimp, marmoset, mouse lemur and bushbaby. In mouse, the homologous region is flanked by ID2 and B4 SINE repeats, whereas it is next to a WALLSI4_Mar SINE in opossum and to a Mon1g1 (MIR family) in platypus. No retrotransposons are seen in the region in the genomes of chicken and xenopus, but in coelacanth there are LFSINE and LmeSINE1b repeats flanking the blast hit. These data suggest that, even though some core sequence of the BRE6 element was present in early vertebrate genomes, it was extended by SINE-mediated transduction probably in early mammals, and acquired its full-length in the genome of early primates.

### 4.2. BRE7

(chr1:87,821,614-87,822,820) is the region originally found in our screening, about 7 kb from the 3’-end of *LMO4*. It shows epigenetic marks of active enhancer in brain germinal matrix and fetal brain according to Roadmap data, and it is located inside an intron of *LINC01364*, a lncRNA with no clear promoter marks and expressed almost exclusively in testis but not in brain. BRE7 overlaps one p300 peak in mouse telencephalon and two Vista elements (element 174 and element 322), both of which show strong enhancer activity in E11.5 mouse forebrain. ENCODE data shows binding of REST (a transcriptional repressor) in neuroectodermal cell line PFSK-1. Hi-C from fetal brain shows increased contacts with *LMO4* promoter, with the TSS of *LINC01364,* and with two other potential brain enhancers (one of them deeply conserved) located at chr1:87857347-87860390 and chr1:87874564-87876915. We searched for expression of this element using blast in RefSeq RNA (all species) but only found two lncRNAs annotated automatically by Gnomo in pig (XR_002343644.1) and in *Parus major* (XR_002002032.1).

COGE blast using this 1.2 kb query found one significant hit in lamprey (E-value= 6e-59) located in scaffold GL492185:3143-3368. Unfortunately, this scaffold is rather short, so no gene is found in the vicinity. There is a high quality hit in elephant shark spanning 700 nt of the query, which is also found in the rest of species and is extended almost completely in the genome of opossum (Figure 7). However, all Osteichthyes (including zebrafish, fugu, spotted gar and coelacanth) show additional hits in different chromosomes/scaffolds, covering nucleotides 680-810 of the query. A more detailed study of microsynteny of the region (see Figure 6) shows that *LMO4* is flanked by *PKN2* and *HS2ST1* in humans and eutherian mammals, with all three genes in the same orientation. In elephant shark, the homologous region to BRE7 is 17.5 kb away from the *LMO4* ortholog, between *HS2ST1* and *PKN1* as in mammals. In fugu, the most significant blast hit (chr20:899,061-899,708) is 13 kb from *LMO4*, syntenic with *HS2ST1* but with no sign of *PKN1/PKN2* on the other side; however, there is a second hit in a different chromosome (chr6:617,238-617,316) 1.6 kb from an ENSEMBL gene which codes for part of human LMO4 protein. In zebrafish, the longest blast hit is 41 kb from *LMO4* ortholog in chr6:25,901,789-25,902,502, with perfect microsynteny. In this species there is also a second blast hit in chr5:32,356,783-32,356,912 located 11 kb from a paralog called *lmo4a*, and an additional third hit in chr8:52,671,535-52,684,434 located near *tubb2*. In spotted gar we found one main blast hit in chrLG10:7419163-7420000, 18 kb away from the *LMO4* ortholog, plus a second hit in a different scaffold (chrLG21:3025062-3030000) between the orthologs for *TUBB4Q* and *C9orf43-POLE3*. Finally, coealancanth also shows a main hit (JH128381:55250-54416) 31 kb from *LMO4* (syntenic with *HS2ST1* on one side, the other side is cut in the assembly), plus a second hit (JH128026:275289-275434) 33 kb from of a paralog called *lmo4.2*. These data indicate the presence of a duplication of this region in the genomes of Actinopterygii and Sarcopterygii that includes a copy of *LMO4* plus the sequence of the BRE6 regulatory element. It is likely that a second duplication took place in the genome of zebrafish, also containing the central core (nucleotides 680-810) of the sequence of this regulatory element.

**Figure 7.**
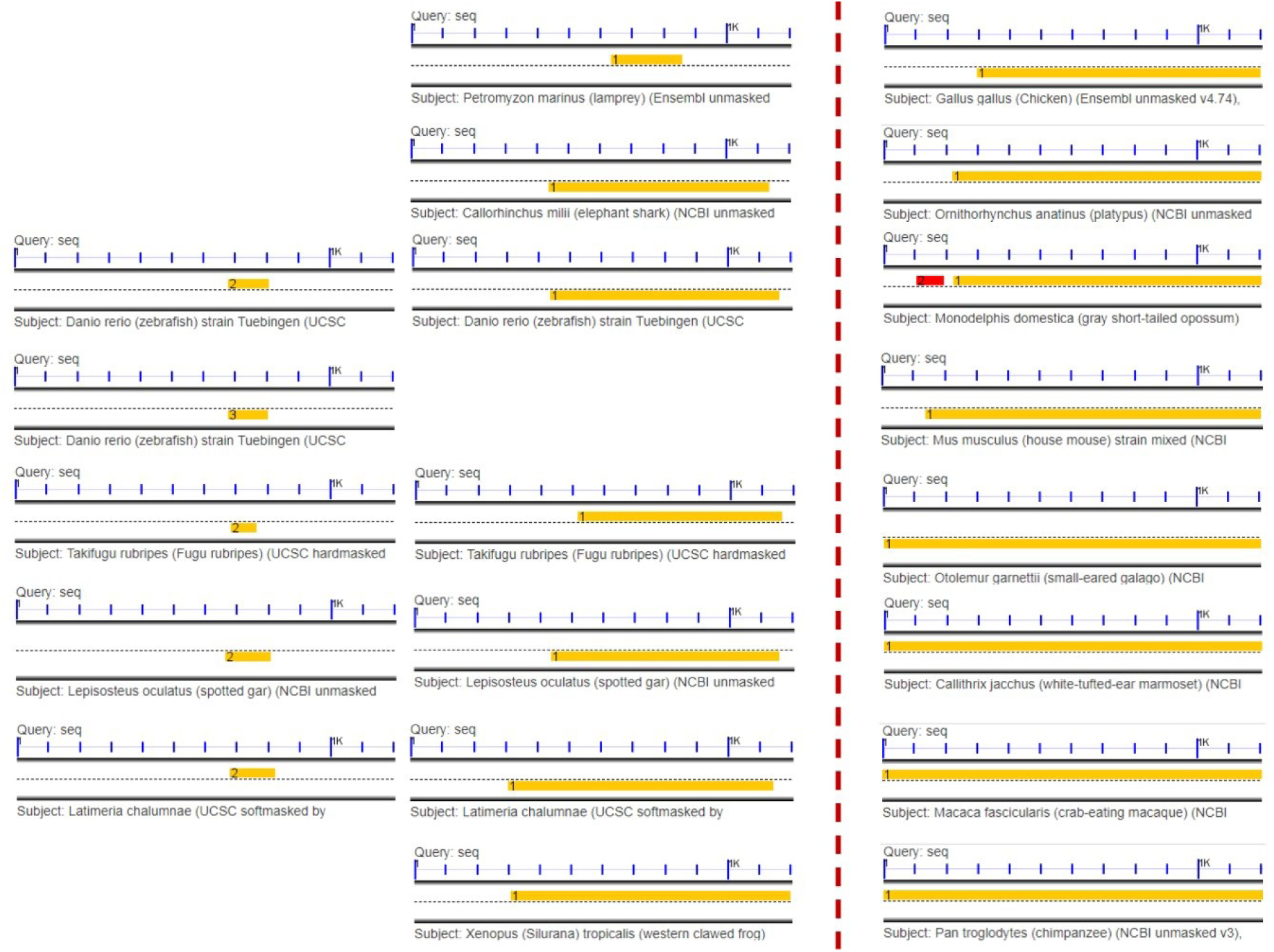
COGE blast results using the human sequence of BRE7 as query. For each hit, a rule at the top represents the size of the query, and a yellow rectangle indicates the size and position of the hit with respect to the query. A hit is found in lamprey which is extended in vertebrates (top to bottom, middle column); extra hits were found in a different chromosome/scaffold in Osteichthyes (shown to the left of zebrafish, fugu, spotted gar and coelacanth). Hits in chicken and mammals are shown to the right of the vertical dashed line (top to bottom).

There is a single blast hit in xenopus (GL172721: 2876588-2877479; assembly JCI 4.2) 30 kb away from *lmo4*, syntenic with *hs2st1* on one side. *Pkn1/2* is also located on the other side, but 1 Mb away from *lmo4* and in the opposite orientation. This suggests that there was an inversion that separated the two genes in this species. Interestingly, xenopus also has the *lmo4a* paralog found in fish, but with no traces of the BRE7 regulatory element nearby.

In chicken, there is one blast hit in chr8:15,776,515-15,777,419 (assembly Galgal4.74) about 10 kb from *LMO4*, with *HS2ST1* on one side and *PKN1* on the other, all in the same orientation. A similar configuration is found in platypus and all other mammals with the exception of opossum. In this marsupial, the blast hit is located in chr2:39,473,771-39,474,873, 17 kb away from *LMO4* with syntenic *HS2ST1* on one side, but the ortholog of *PKN1/2* is located 23 Mb away from *LMO4* on the same side of *HS2ST1* and in the opposite orientation (see Figure 6 for details). This suggests an inversion in the genome of this species that took apart *PKN1/2* and *LMO4* but did not disrupt the association of these two genes with BRE6 or BRE7, respectively.

Interestingly, BRE7 is flanked by a MIR3 SINE repeat on one side and an L2a LINE on the other in human and also in the genomes of all other primates analysed from chimp to galago (*Otolemur garnetii)*. The same happens in mouse, although there are two additional SINEs (MER131 and B4A) in the middle of BRE7. Likewise, the homologous region is flanked by MIR3_MarsA and L2_Mars in opossum. In platypus, it is flanked by Mon1a1 (family MIR) and L2_Plat1o repeats, with some MER family elements inside the region. In chicken, the homologous BRE7 region is flanked by MER131 SINE repeats, but in the genome of xenopus it is flanked by a HAT-Charlie DNA transposon and a Gypsy LTR. No recognizable transposable elements are found in the region in coelacanth, elephant shark and lamprey genomes.

In summary, the sequence corresponding to the BRE7 regulatory element appeared near *LMO4* very early in gnatosthome evolution coinciding with the appearance of the gene, and acquired its full length in early mammals probably via SINE-mediated transduction. Duplications of *LMO4* orthologs in Osteichthyes kept the association with the regulatory element. Likewise, the genomes of xenopus and opossum underwent inversions that broke the synteny of the region but did not affect the association of the element with *LMO4* orthologs.

### 4.3. BRE8

(chr1:88,758,718-88,759,914) is included in a strong peak of Hi-C contacts with the promoter of *LMO4* in fetal brain cortical plate and germinal zone (Supplementary Figure S6), even though it is located 950 kb away. It does not overlap any Vista enhancer, FANTOM5 enhancer or p300 peak, but it has good Roadmap marks of active enhancer exclusively in brain tissues. ENCODE data shows binding of three TFs (TCF7L2, STAT1 and ESR1) in some cell lines. About 10 kb downstream from this site there is another fragment with epigenetic marks of active enhancer in brain tissues that binds CTCF, SMC3 and RAD21, so it probably represents an insulator that could acts as boundary for a TAD. Although BRE6 is closer to *PKN2* than to *LMO4*, Hi-C data suggest that it regulates the expression of *LMO4* at least in human fetal brain.

COGE blast with the human sequence of BRE6 found significant hits only in mammals, with a fragment of 300 bp of the query present in platypus, opossum and mouse that is extended almost completely in *Otolemur* and the other primate genomes studied. The binding sites for the three TFs mentioned above lie within this primate-specific fragment. In opossum, the element is separated from the *PKN2* ortholog by the inversion mentioned previously (Figure 6). Interestingly, we found a EULOR7 transposon (Eutelostomi low-copy repeat) next to the sequence of the BRE6 element from platypus to primates, so this element probably originated via a transposon-mediated transduction event in early mammals. Analysis of expression using blast on RefSeq RNA was negative.

## DISCUSSION

In recent years, several lines of research have improved our understanding of chromatin function in the context of 3D conformation. The initial view of promoter-enhancer interactions, according to which a regulatory element would regulate the activity of the closest gene, has been gradually replaced by the loop extrusion model in which the main constraint of promoter-enhancer interaction is their presence in the same TAD, regardless of the distance between them (Dekker and Mirny 2016). In fact, several recent experimental studies analysing promoter-enhancer interactions have confirmed that the gene regulated by an enhancer is not necessarily the closest one (Shlyueva et al. 2014; Smemo et al. 2014; Fulco et al. 2016). In the present work, we have devised an analysis pipeline starting with genomic regions previously predicted as enhancer elements in high-throughput projects and then integrates information from biological datasets indicative of (i) enhancer activity (open chromatin, epigenetic marks, TF binding), and (ii) physical contact of putative regulatory regions with the promoters of genes in the vicinity. A similar pipeline was recently used with high predictive success by Fulco et al. in the analysis of enhancer elements regulating the expression of *MYC* and *GATA1* (Fulco et al. 2016). In our study, the availability of different datasets from similar tissues, brain regions and developmental stages enabled us to make good predictions not only about the activity of a genomic region as regulatory element, but also about the gene(s) most likely to be regulated. A case in point is the element identified here as BRE1: using p300 ChIP-seq in dorsal cerebral wall of E14.5 mice, Wenger et al. identified an enhancer close to the promoter of *Tbr1* (Wenger et al. 2013), but failed to link the regulation of this gene with a more distal element corresponding to BRE1; analysis of Hi-C datasets allowed us to unambiguously establish physical contact between them, at least in human brain. Although promoter-enhancer contact is not definitive proof of functional causality, Fulco et al. found that quantitative measures of chromatin state and chromosome conformation are strongly predictive of enhancer functionality, correctly ranking 6 out of 7 distal *MYC* enhancers in their study. We also note that not all regions showing high contact frequency with a promoter (by Hi-C) displayed strong enhancer activity (by epigenetic marks and chromHMM segmentation analysis); this is probably due to the fact that the source of samples was not identical: Hi-C data came from fetal cortical plate and germinal zone, whereas chromatin epigenetic marks were analysed in several regions of adult brains or in entire fetal brain samples.

We have focused on elements that very likely regulate the expression of three genes involved in neocortex development (Hermanson et al. 1999; Cecchi et al. 2000; Bulchand et al. 2003; Rakic 2009; Mariani et al. 2012; Wenger et al. 2013). BRE1 is a deeply conserved element that might upregulate *TBR1* expression in pallium in vertebrates. Interestingly, *Tbr1* is known to be expressed in chicken, mouse and zebrafish brain, but we could not find information about expression in elephant shark or lamprey. Given that this regulatory element is not present in lamprey but both lamprey and elephant shark have a *TBR1* ortholog, it would be interesting to analyse the expression of this gene in the pallium of these two species. This would be particularly interesting in the case of lamprey, as lamprey pallium shows marked similarities with mammalian neocortex in terms of efferent connectivity (Ocaña et al. 2015).

The regulatory landscape of *EMX2* shows one deeply conserved element (present in elephant shark) close to the gene, two more elements that seem mammalian innovations (one appearing in opossum and the other in mouse) and another distal element with strong Hi-C contact peaks that is a clear primate innovation. As this gene is important in forebrain regionalization and archipallium development in mammals, it will be interesting to study the effect of these regulatory elements on *EMX2* expression in platypus, opossum, mouse and primates.

A similar regulatory landscape was observed in the case of *LMO4*, with one deeply conserved element present in lamprey and close to the gene, a second element appearing in coelacanth and a third one appearing in platypus that looks a mammalian innovation. *Lmo4* expression in mouse suggests involvement in the development of some parts of telencephalon (Hermanson et al. 1999; Bulchand et al. 2003); in zebrafish, *lmo4b* (the ortholog of human *LMO4*) is also expressed in the neural plate and telencephalon, and it seems to limit the size of the forebrain through inhibition of *six3b* and *rx3* (Lane et al. 2002; McCollum et al. 2007). In summary, the general picture obtained from the study of CNEs of these three genes shows a mosaic of ancient and more recent regulatory elements, consistent with deep vertebrate homology for some CNEs (McEwen et al. 2009) upon which mammal and primate innovations accrued. It will be interesting to study the correlation of these regulatory landscapes with the evolution of gene expression and telencephalon development.

The evolutionary analysis of the eight regulatory elements included in our study sheds some light on their origin, either by duplication or by transposon-mediated transduction. MacEwen et al. have shown that duplication was an important driver of the expansion of ancient vertebrate CNEs (McEwen et al. 2009); this is what we found for BER1, a regulatory element concurrent with the duplication of *AZI2* that created *TANK* during the transition from agnatha to gnathosthomes. Interestingly, this duplication is accompanied by an inversion of upstream gene *RBMS1*; this rearrangement could have originated BRE1 by bringing some sequence into the proximity of the newly created copy of *AZI2* as it was being neofunctionalised into *TANK*. Notwell et al. (Notwell et al. 2015) have previously found enrichment of MER130 elements in mouse brain developmental enhancers, suggesting that they contributed TF binding sites. We have not seen transposable elements overlapping binding sites for TFs in the regulatory elements studied in this work, but at least five of them (BRE2, BRE4, BRE5, BRE6 y BRE8) are flanked by SINE family members, suggesting their evolution by some type of transposon-mediated transduction event.

We have devised an analysis pipeline that integrates available datasets in order to identify and characterize regulatory elements involved in brain development and evolution. However, a complete picture of how the evolution of genomic regulatory landscapes has driven the evolution of the primate brain will require the integration of detailed transcriptomic, epigenomic, TF-binding and 3D chromatin conformation data from homologous brain regions of different mammals and non-mammalian vertebrates at several developmental stages. Although this might seem a distant target, recent comparative transcriptomic studies point in that direction (Bakken et al. 2016; He et al. 2017) and we can only hope that future studies will advance towards that goal. In the meantime, our work could help to guide experimental studies in mammalian and non-mammalian vertebrate models aimed at elucidating the role of conserved regulatory elements in brain evolution.

## METHODS

### Tools for the identification of regulatory elements

We downloaded a set of predefined enhancers from the FANTOM5 project (Andersson et al. 2014) selecting enhancers specific for astrocytes, neuron, neural stem cells (under cell-specific enhancer category) and for brain (organ-specific enhancers) from (http://slidebase.binf.ku.dk/human_enhancers/presets). This dataset comprised 2,019 enhancers (associated to 2,597 genes according to analysis performed with GREAT) which were uploaded as a custom track to the UCSC Genome Browser (human assembly hGRC37/hg19).

We used several datasets from the Roadmap Epigenomics Project (Kundaje et al. 2015) available at the UCSC Genome Browser as a track hub (“Human Epigenome Atlas Data Complete Collection, VizHub at Washington University in St. Louis”). In order to predict open chromatin regions, we used DNAse hypersensitive sites (DHS) tracks in the following neural cells and tissues: E054 Ganglion Eminence derived primary cultured neurospheres; E053 Cortex derived primary cultured neurospheres; E068 Brain Anterior Caudate; E069 Brain Cingulate Gyrus; E071 Brain Hippocampus Middle; E072 Brain Inferior Temporal Lobe; E073 Brain Dorsolateral Prefrontal Cortex; E074 Brain Substantia Nigra.

For chromatin segmentation we used the tracks generated by chromHMM (Ernst and Kellis 2012) for the following neural and non-neural tissues and cells: E003 H1 Cells; E082 Fetal Brain Female; E081 Fetal Brain Male; E108 Skeletal Muscle Female; E107 Skeletal Muscle Male; E066 Liver; E101 Rectal Mucosa Donor 29; E102 Rectal Mucosa Donor 31; E062 Primary mononuclear cells from peripheral blood; E033 Primary T cells from cord blood; E034 Primary T cells from peripheral blood; E038 Primary T helper naive cells from peripheral blood; E031 Primary B cells from cord blood; E032 Primary B cells from peripheral blood; E068 Brain Anterior Caudate; E069 Brain Cingulate Gyrus; E071 Brain Hippocampus Middle; E072 Brain Inferior Temporal Lobe; E073 Brain Dorsolateral Prefrontal Cortex; E074 Brain Substantia Nigra; E116 GM12878 Lymphoblastoid Cells; E122 HUVEC Umbilical Vein Endothelial Primary Cells; E110 Stomach Mucosa; E070 Brain Germinal Matrix; E067 Brain Angular Gyrus; E113 Spleen; E098 Pancreas.

In order to find transcription factor binding sites (TFBS), we accessed ChIP-seq data available at the UCSC Genome Browser that were processed using the computational pipeline developed by the ENCODE Project (Bernstein et al. 2012) to generate uniform peaks of TF binding. In this dataset, peaks for 161 transcription factors in 91 cell types are combined into clusters to produce a display showing occupancy regions for each factor and motif sites (Wang et al. 2012; Gerstein et al. 2012).

We used information about Vista enhancers available as a track hub from the UCSC Genome Browser. Vista enhancers are non-coding elements identified as conserved in human, mouse, and rat; each element was cloned into a reporter vector with a minimal promoter fused to LacZ, injected into fertilized mouse eggs and 11.5 day embryos were assayed for lacZ activity (Pennacchio et al. 2006).

Candidate enhancer regions in mouse embryonic telencephalon were obtained from Wenger et al. (Wenger et al. 2013) who used ChIP-seq to detect p300 binding in the genome of mouse E14.5 dorsal cerebral wall samples. We downloaded 6,340 peaks of p300 binding in mouse (mm9 assembly), converted them to human hGRC37/hg19 coordinates with liftover and uploaded as a custom track to the Genome Browser.

### Tools for analysis of gene expression and long-range chromatin interactions

Gene expression data were obtained from the GTEx Consortium (Melé et al. 2015) and from Brainspan (Miller et al. 2014). Data from Brainspan for specific genes were viewed in GenTree (http://gentree.ioz.ac.cn/index.php).

In order to find support for enhancer-promoter contacts we used functional information about eQTLs (expression Quantitative Trait Loci) available from GTEx; likewise, we searched for single nucleotide polymorphisms (SNPs) identified by published Genome-Wide Association Studies (GWAS) in the vicinity of candidate regulatory elements, using the track available at the UCSC Genome Browser.

Physical interactions between candidate regulatory elements and surrounding gene promoters (up to 1 Mb in either direction) ware assessed inspecting Hi-C datasets from various cells/tissues, particularly those from human fetal cortical plate and germinal zone (Won et al. 2016). Hi-C data were viewed as virtual 4C plots in the 3D Genome Browser (Wang et al. 2017); as they were not normalized to correct for the decay of signal as a function of the distance, distant peaks are more likely to represent true contacts. Likewise, performing “reverse” 4C plots (using either the enhancer or the promoter as anchoring point) helped to identify contacs with high confidence.

### Tools for conservation analysis

We accessed 481 ultraconserved segments longer than 200 bp that are absolutely conserved between orthologous regions of the human, rat, and mouse genomes (Bejerano 2004) using the track hub available at the UCSC Genome Browser. We also downloaded a set of 36,857 ancient CNEEs (conserved non-exonic elements) described by Lowe et al. (Lowe et al. 2011) and we uploaded this dataset as a custom track in the UCSC Genome Browser.

Alignment of candidate regulatory regions with genomic sequences was performed with Blast (Altschul et al. 1990) implemented either at the NCBI (https://blast.ncbi.nlm.nih.gov/Blast.cgi) or as part of the CoGe suite (Lyons and Freeling 2008) that was originally developed to compare plant genomes (https://genomevolution.org/coge/). Each regulatory element was inspected in detail, selecting the core region more likely responsible for enhancer activity (open chromatin, epigenetic marks, TF binding). We obtained the human sequence (repeat-masked when appropriate) and used this as query in blastn searches against genomes from several species representing a wide range of phyla from sponges to primates in the COGE platform. We used an E-value cut-off of 10^−6^ to identify homologous regions, which is standard practice for blastn searches. Since initial searches only revealed significant hits in vertebrate genomes, we focussed our analysis on vertebrates using representative genomes from lamprey to chimp (see Table 2 for genome assemblies used).

**Table 2.**
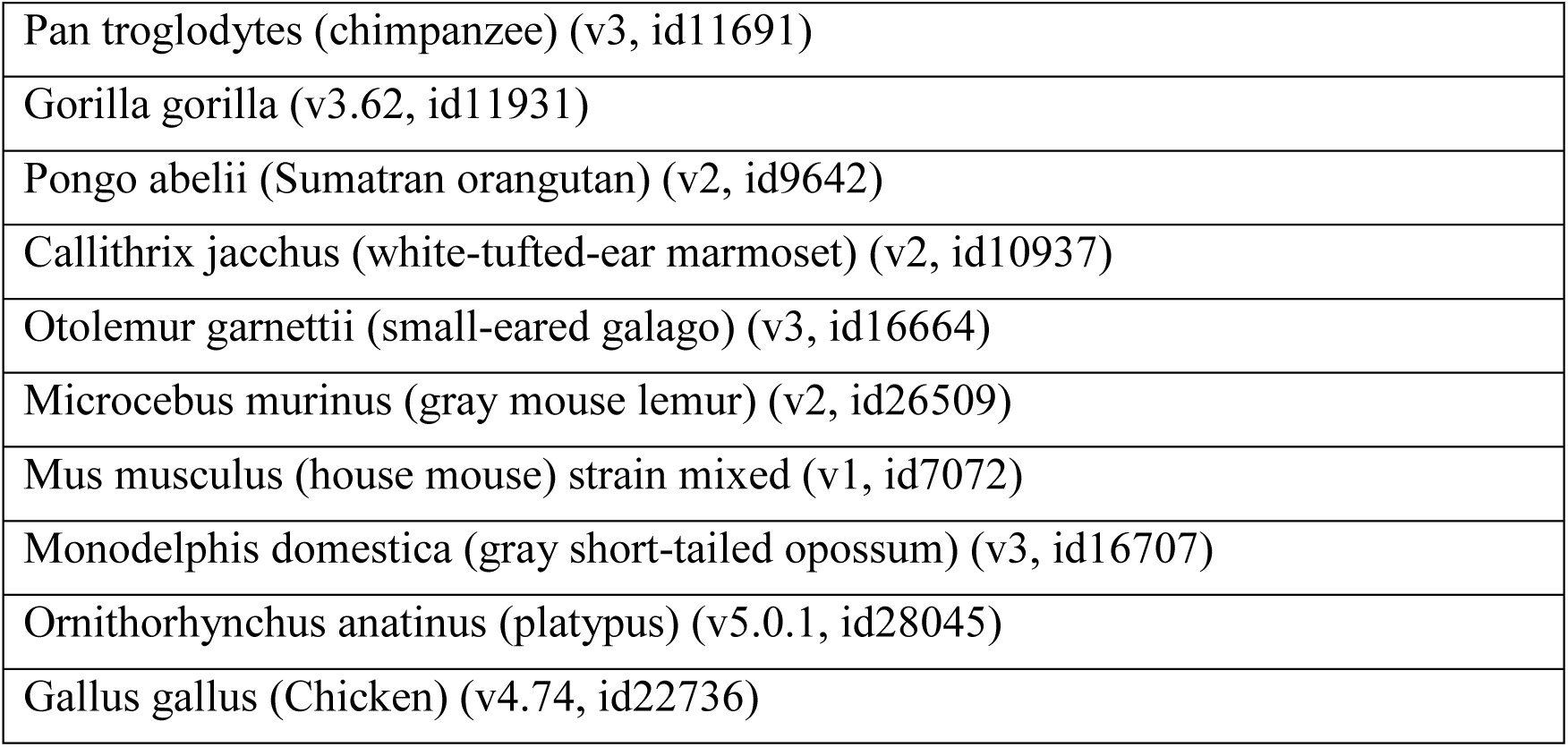
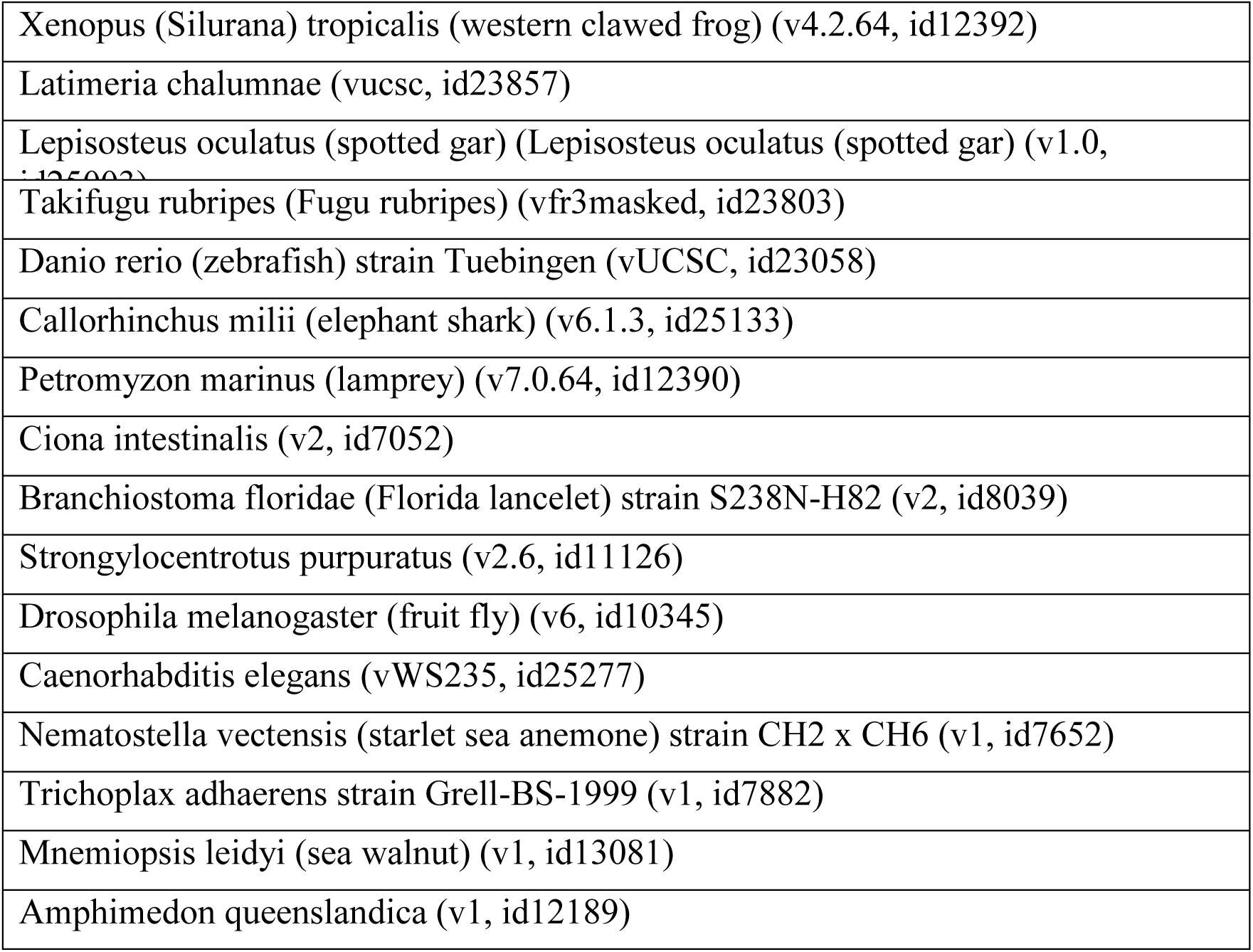
Genomes and assemblies used in COGE blastn searches.

In order to check whether enhancer sequences were expressed (or part of expressed sequences) we used blastn at NCBI Blast with the same queries on EST and RefSeq RNA sequences (refseq_rna database). Multiple alignments of some genomic regions were performed with Shuffle-LAGAN (Brudno et al. 2003) at the VISTA suite (http://genome.lbl.gov/vista/index.shtml).

## ACKNOWLEDGEMENTS

I wish to thank all departmental colleagues and students with whom I have engaged in helpful discussions about this work, in particular José L. Vizmanos.

## DECLARATIONS

The author declares no potential competing interests in the manuscript.

## ADDITIONAL INFORMATION

**Figure S1.**
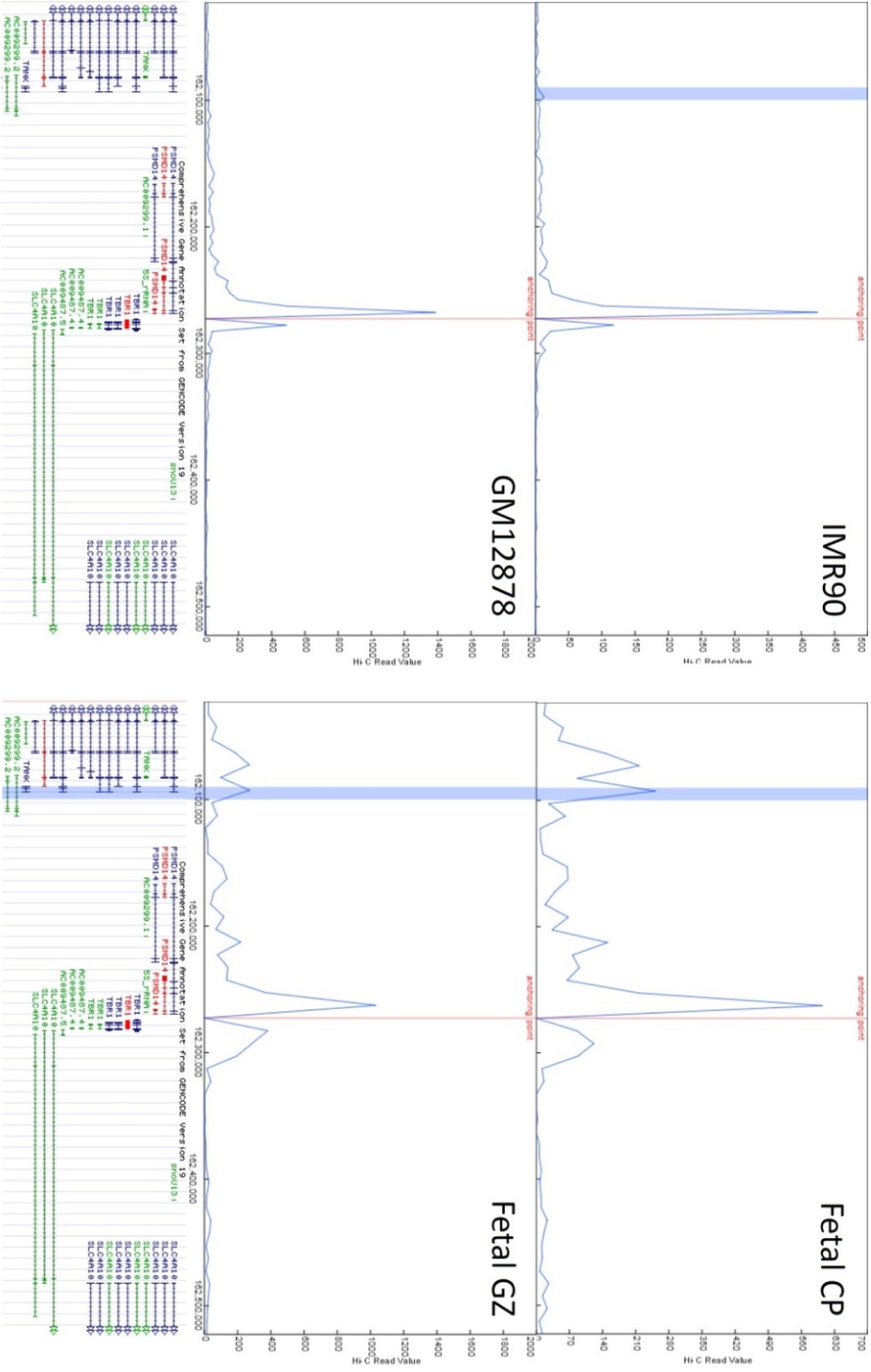
Virtual 4C plots using Hi-C data in non-brain cell liness (GM12878 and IMR90), fetal brain cortical plate (Fetal CP) and germinal zone (Fetal GZ). In each plot, the anchoring point (red line) marks the promoter of *TBR1*. There is a peak of contacts with BRE1 (region highlighted in blue at the 3’-end of *TANK* transcripts) only in brain samples. The bottom panel shows the GENCODE gene annotation comprehensive set, version 19. Contacts (Y-axis) are shown as RPKMs.

**Figure S2.**
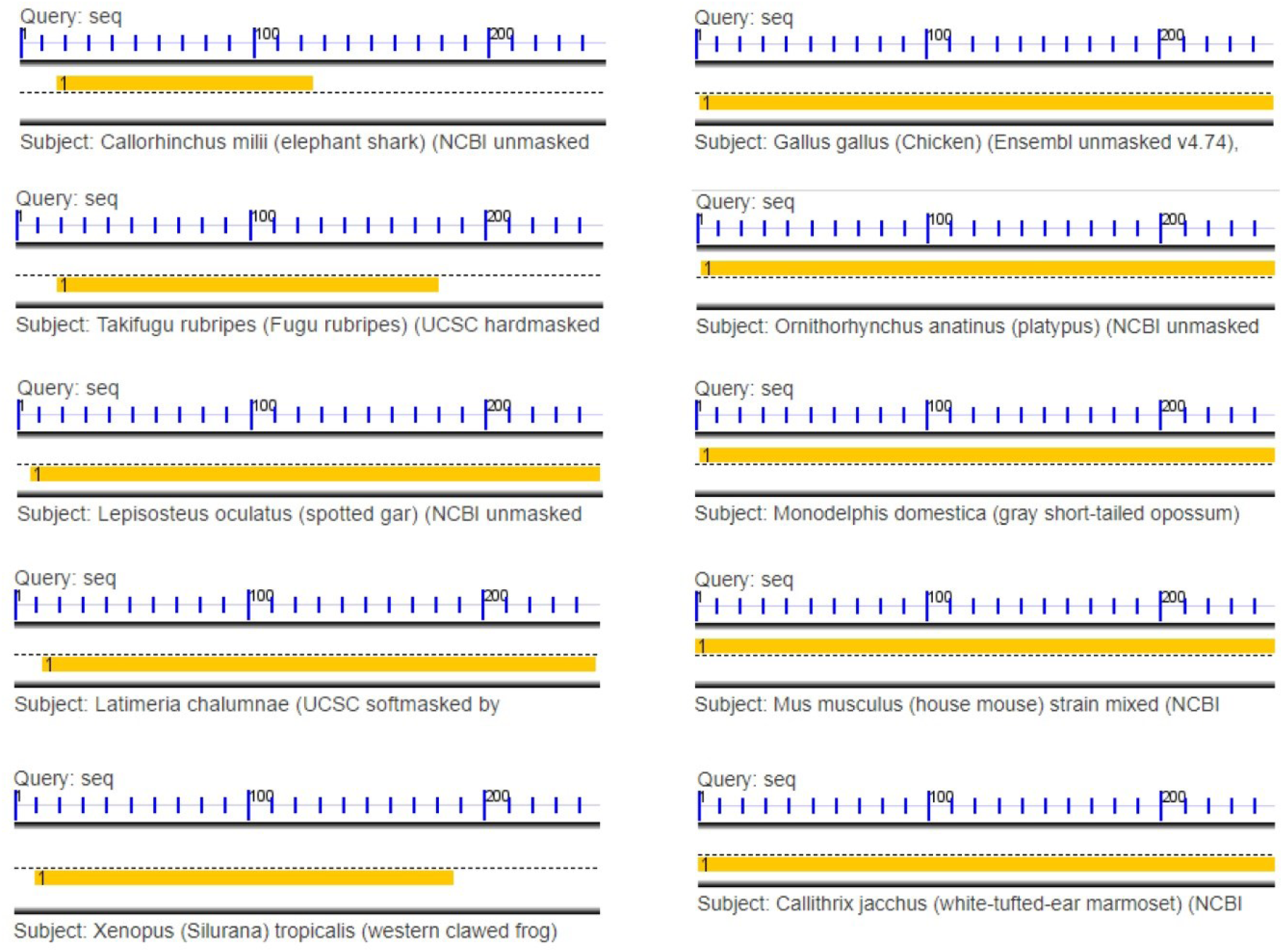
COGE blast results using the human sequence of BRE1 as query. For each hit, a rule at the top represents the size of the query, and a yellow rectangle indicates the size and position of the hit with respect to the query. The oldest hit is found in elephant shark and it expands in later vertebrates (top to bottom, left column first). From chicken to primates, the hit covers the entire query.

**Figure S3.**
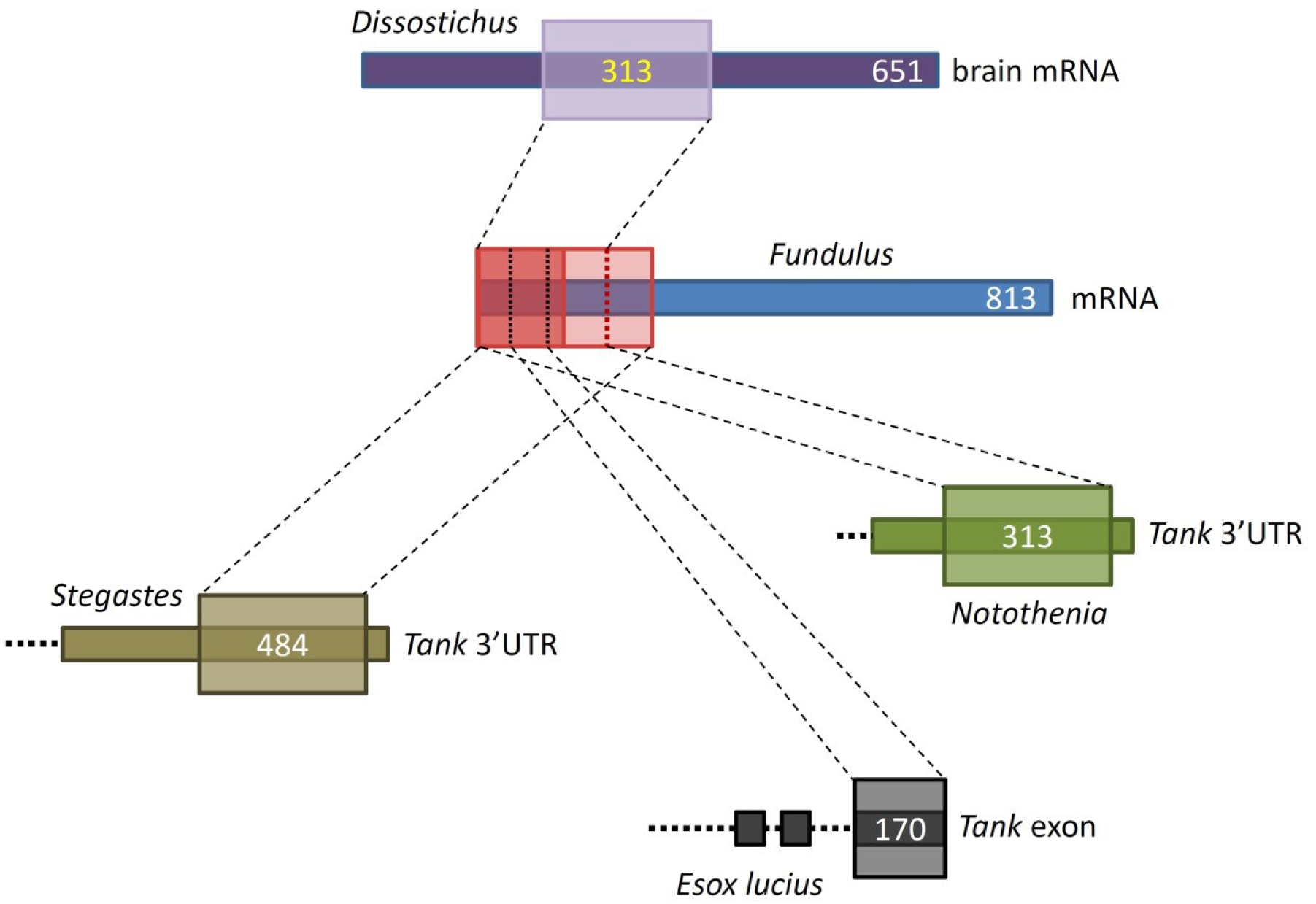
Schematic representation of blast hits for the BRE1 sequence in expressed sequences from EST and RefSeq RNA databases. The whole query is present in an mRNA from *Fundulus grandis* (red rectangle); parts of the query also find significant hits in expressed sequences from other Acanthopterygii, frequently as part of the 3’-end of *Tank* mRNAs (see text for details).

**Figure S4.**
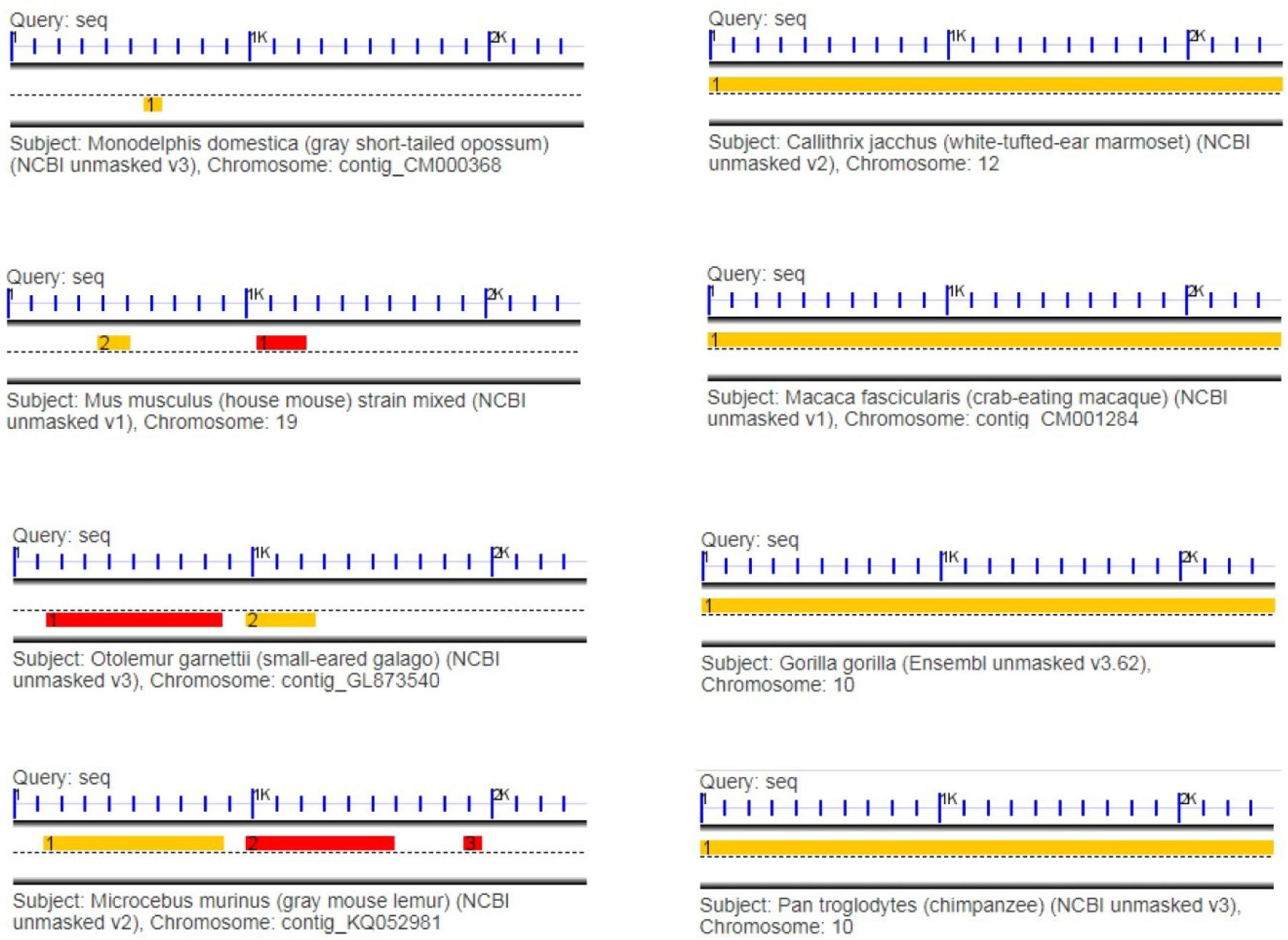
COGE blast results using the human sequence of BRE2 as query (top to bottom, left column first). For each hit, a rule at the top represents the size of the query, and a yellow rectangle indicates the size and position of the hit with respect to the query. The oldest significant hit is found in opossum (top left) and a second hit appears in mouse; both hits are progressively extended in primates.

**Figure S5.**
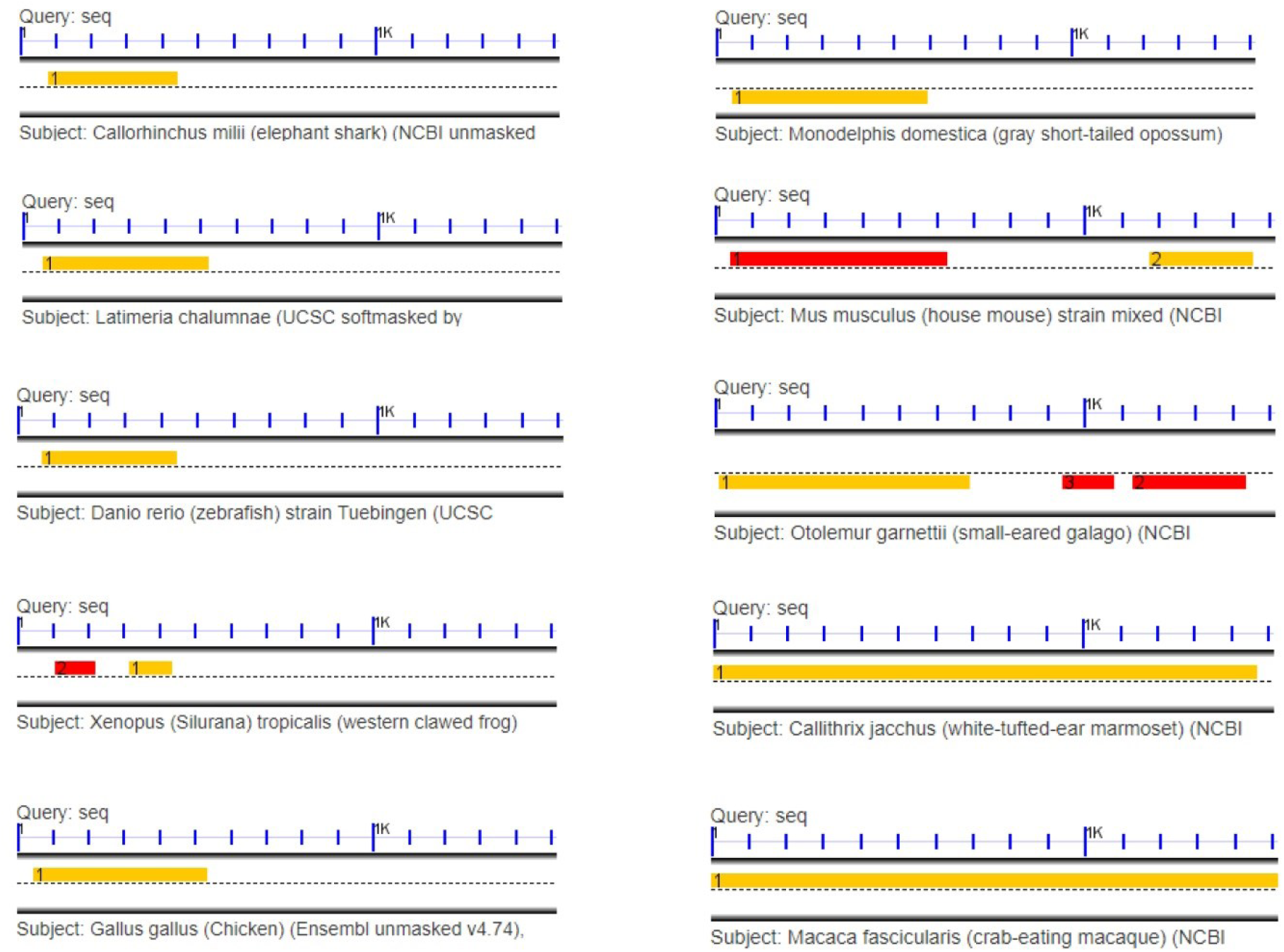
COGE blast results using the human sequence of BRE3 as query (top to bottom, left column first). For each hit, a rule at the top represents the size of the query, and a yellow rectangle indicates the size and position of the hit with respect to the query. The oldest hit is found in elephant shark and it is conserved with the same size until chicken (with the exception of xenopus, where it is broken into two fragments). A second hit appears in mouse and both hits are progressively extended in primates.

**Figure S6.**
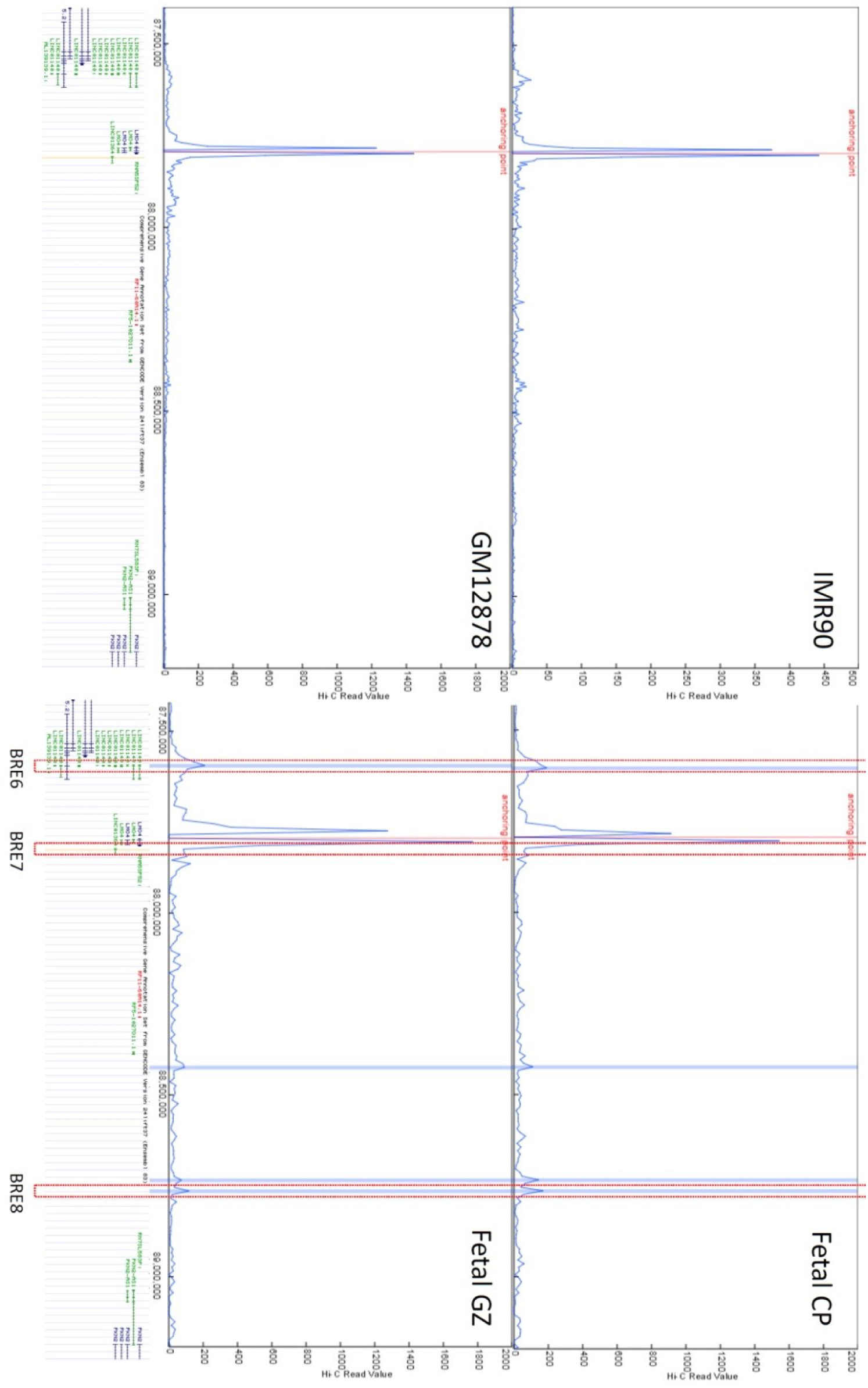
Virtual 4C plots using Hi-C data from non-brain cell lines (GM12878 and IMR90), fetal brain cortical plate (Fetal CP) and germinal zone (Fetal GZ). In each plot, the anchoring point (red line) marks the promoter of *LMO4*. Several peaks of contacts indicate the location of BRE6, BRE7 and BRE8 (highlighted by red rectangles), active in brain samples but not in GM12878 or IMR90 cells (note the different scale of the Y-axis in IMR90). The bottom panel shows the comprehensive gene annotation from GENCODE version 24lift37. Contacts (Y-axis) are shown as RPKMs.

